# Multi-cancer analysis of clonality and the timing of systemic spread in paired primary tumors and metastases

**DOI:** 10.1101/825240

**Authors:** Zheng Hu, Zan Li, Zhicheng Ma, Christina Curtis

**Affiliations:** Department of Medicine, Division of Oncology, Stanford University School of Medicine, Stanford, California, USA; Department of Genetics, Stanford University School of Medicine, Stanford, California, USA; Stanford Cancer Institute, Stanford University School of Medicine, Stanford, California, USA; Life Science Research Center, Core Research Facilities, Southern University of Science and Technology, Shenzhen, Guangdong, China

## Abstract

Metastasis is the primary cause of cancer-related deaths, but the natural history, clonal evolution and impact of treatment are poorly understood. We analyzed exome sequencing data from 457 paired primary tumor and metastatic samples from 136 breast, colorectal and lung cancer patients, including untreated (n=99) and treated (n=100) metastatic tumors. Treated metastases often harbored private ‘driver’ mutations whereas untreated metastases did not, suggesting that treatment promotes clonal evolution. Polyclonal seeding was common in untreated lymph node metastases (n=17/29, 59%) and distant metastases (n=20/70, 29%), but less frequent in treated distant metastases (n=9/94, 10%). The low number of metastasis-private clonal mutations is consistent with early metastatic seeding, which we estimated commonly occurred 2-4 years prior to diagnosis across these cancers. Further, these data suggest that the natural course of metastasis is selectively relaxed relative to early tumor development and that metastasis-private mutations are not drivers of cancer spread but instead associated with drug resistance.

## Introduction

Metastasis remains poorly understood despite its critical clinical importance. For instance, metastases have been reported to originate from a single cell or clone in the primary tumor (monoclonal seeding) ^1-4^ or multiple clones (polyclonal seeding) ^5-7^, but the prevalence of these patterns across distinct tumor types is unknown as is the impact of therapy and the timing of metastatic seeding ^8-10^. While several recent studies have genomically characterized metastatic lesions in the absence of the matched primary tumor ^11-13^, with such data it is not feasible to disentangle the drivers of metastasis from those that are treatment associated since metastases are often sampled after treatment. However, comparisons of paired primary tumors and metastases have been far more limited due to the challenge in obtaining such samples ^5,8,14-18^. As such, there has yet to be a systematic analysis of monoclonal versus polyclonal seeding, the chronology of systemic spread and the effect of therapy across cancers.

Here we analyzed whole-exome sequencing (WES) data from 457 paired primary tumor (P) and metastases (M) from 136 patients with colorectal, lung or breast cancers using a uniform bioinformatics pipeline. We assessed ‘driver’ gene heterogeneity and evaluated the prevalence of monoclonal versus polyclonal seeding, revealing considerable variability between untreated and treated metastases across cancer types. Treatment was associated with high primary tumor versus metastasis (P/M) driver gene heterogeneity and monoclonal metastases. Metastatic seeding was estimated to occur two to four years prior to diagnosis of the primary tumor across three common cancer types, with breast cancers generally disseminating later and therefore closer to the time of detection relative to colorectal and lung cancers. Collectively, these observations suggest that systemic spread can begin early during tumor growth and that clonal architecture is remodeled by treatment, providing new insights into the clonal evolution of metastasis.

## Results

### The landscape of genomic alterations in paired primary tumors and metastases

We performed a literature review to identify cohorts with genomic sequencing data from matched normals, primary tumors (P) and metastases (M) from patients with three common cancer types, namely, colorectal^16,17,19-22^, lung^23,24^ and breast^23,25-29^ (**Tables S1-S2, Fig. S1**). All samples were processed within a uniform bioinformatics pipeline ^16,30^ to identify somatic single nucleotide variants (SSNVs), insertions/deletions (indels) and somatic copy number alterations (SCNAs) (**Methods**). Tumor purity/ploidy and cancer cell fraction (CCF) of SSNVs and indels (referred as SSNVs hereafter) were estimated in order to distinguish clonal (the upper bound of 95% confidence interval or CI of CCF ≥ 1) versus subclonal (the upper bound of 95% CI of CCF < 1) SSNVs (**Methods**). Following quality control assessment (**Methods**), we retained 457 tumor samples from 136 patients (colorectal cancer, n=39; lung cancer, n=30; breast cancer, n=67) for downstream analysis (**Tables S1-S3, Fig. S1**).

Metastases exhibited higher purity than paired primary tumors while ploidy was comparable between primary tumors and metastases in all three cancer types (**Fig. S2**). Overall, the mutational burden (SSNVs or SCNAs) was highly concordant between P/M pairs (**Figs. S3-5, Table S4**), although differences between cancer types were noted. For instance, in breast cancer, the SCNA burden between P/M pairs was more concordant than the SSNV burden (**Fig. S3**), a pattern that is more evident in ER+/HER-subgroup (**Fig. S5**). Although primary breast cancers can be copy number driven ^31,32^, these data suggest that metastases can acquire substantial SSNVs and this seemed especially true of ER+/HER-breast cancers, which are often exposed to endocrine therapy and tend to recur later ^33^. In all three cancer types, metastases exhibited a slight increase in the number of clonal SSNVs and fewer subclonal SSNVs (**Fig. S4**), consistent with an evolutionary bottleneck during metastasis. The mutational spectrum of M-private SSNVs (clonal or subclonal) between treated and untreated metastases was also highly concordant except that treated colorectal metastases were characterized by an enrichment of T>G transversions relative to untreated samples (**Fig. S6**). Indeed, all treated colorectal metastases (n=7) were biopsied after 5-fluorouracil (5-FU) chemotherapy in this cohort, which was recently shown to be associated with this mutational pattern ^34,35^.

We next evaluated the enrichment of functional driver gene mutations in paired primary tumors and metastases. Three methods, namely PolyPhen-2 ^36^, FATHMM-XF ^37^ and CHASMplus ^38^, were employed to assess the functionality (“driverness”) of nonsynonymous SSNVs in putative driver genes according to TCGA and COSMIC (**Methods, Table S5**). In total, 1085 functional driver SSNVs/indels were detected across these three cancer types (**Fig. 1a-b, Table S6**), in which 733 were clonal (including shared clonal, P-private clonal or M-private clonal) and 352 were subclonal (shared subclonal, P subclonal/M clonal, P-private subclonal or M-private subclonal). Notably, 84%, 86% and 59% of clonal drivers in each P and M were shared in colorectal, lung and breast cancer, respectively, while the fractions of subclonal drivers was 20%, 50% and 23%, respectively (**Fig. 1c**). Of note, colorectal cancer had highest prevalence of P-private subclonal drivers likely because multi-region sequencing (MRS) data was more prevalent for this tumor type (36%, or 14/39) as compared to lung (0%, 0/30) and breast (9% or 6/67) cancer, and MRS increases the power to detect subclonal mutations. Amongst all driver mutations, M-private clonal and subclonal driver mutations were significantly enriched in breast cancer than colorectal and lung cancers (**Fig. 1c**). Gene ontology (GO) analysis of M-private driver genes revealed enrichment for chromatin binding, modification and organization genes (**Fig. S7, Table S7**), implicating chromatin regulators in metastatic progression ^39^.

**Figure 1.**
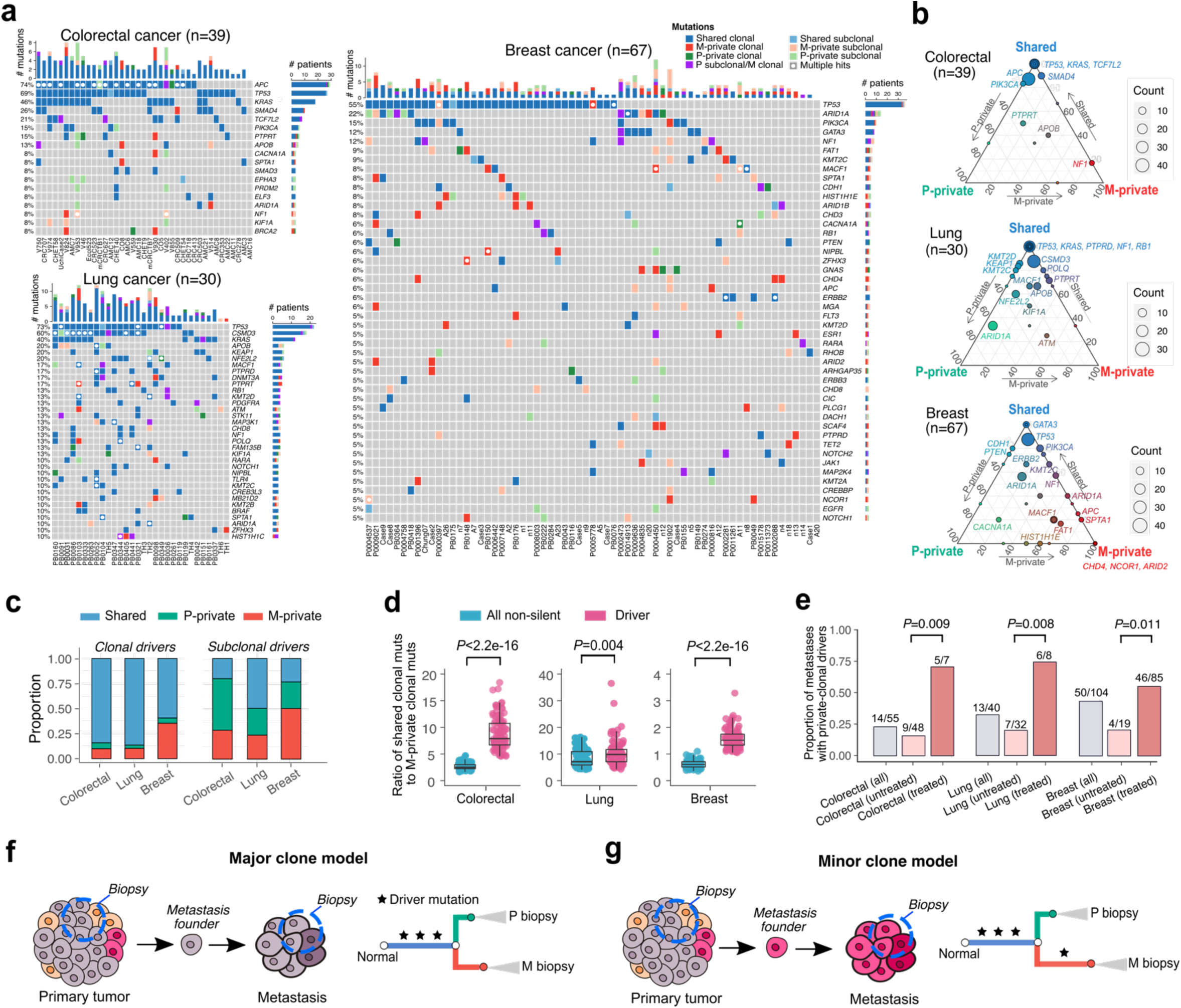
Landscape of driver mutations in paired primary tumors (P) and metastases (M). (**a**) Oncoprint of functional driver mutations in the three cancer types grouped by P/M shared, P-private or M-private mutations including both clonal or subclonal drivers. Genes mutated in at least three patients are shown. Boxes with white circles indicate genes with multiple mutations in a given patient usually in tumor suppressor genes (*TP53, APC, CSMD3*, etc.). (**b**) Ternary plot of mutation counts in driver genes, comparing P-private (left, green), M-private (right, red), and shared (top, blue). The color of each circle indicates the relative frequency of driver mutations among these groups, while the size of the circle represents their overall count in the corresponding cancer type. (**c**) The proportion of shared, P-private or M-private drivers (*clonal*: shared clonal, P-private clonal or M-private clonal; *subclonal*: shared subclonal, P subclonal/M clonal, P-private subclonal or M-private subclonal) in each of the three cancer types. (**d**) The ratio of shared clonal to M-private clonal mutations for all non-silent and driver mutations, respectively. A down-sampling procedure was performed to derive the ratio (Methods) where n=100 down-samplings (50% patients each) were repeated for each of the three cancer types. *P*-value, Wilcoxon Rank-Sum Test (two-sided). Bar, median; box, 25th to 75th percentile (interquartile range, IQR); vertical line, data within 1.5 times the IQR. (**e**) The proportion of metastases harboring at least one private clonal driver mutation grouped by all metastases, untreated and treated metastases. *P*-value, Fisher’s exact test (two-sided). (**f**) Schematic representation of the major clone model where metastasis originates from the major driver clone in the primary tumor leading to driver gene homogeneity between paired P and M biopsies. (**g**) Schematic representation of the minor clone model in which metastases originate from a minor clone in the primary tumor. Due to the inability to detect the minor driver clone in bulk sequencing data, the minor clone model leads to driver heterogeneity between P and M biopsies.

Amongst all non-silent clonal SSNVs in metastases, functional driver mutations were highly enriched on the trunk (P/M shared clonal) of the phylogenetic tree in both colorectal and breast cancers (**Fig. 1d, Methods**). However, this pattern was much weaker in lung cancer (**Fig. 1d**), presumably due to the large number of tobacco-associated non-silent clonal SSNVs (C>A mutations) induced early during lung cancer development (**Fig. S8**) as most of the lung cancer patients in this cohort (∼90%) had a smoking history ^23,24^, whereas driver mutations did not increase proportionally (**Fig. 1d**). In line with these results, the decreased ratio of nonsynonymous versus synonymous SSNVs (dN/dS) ^40^ amongst putative driver genes in metastases (**Fig. S9**) suggests relaxed selective pressure relative to early cancer development in colorectal and breast cancers, but not lung cancer. Only 25%, 33% and 48% of colorectal, lung and breast cancer metastases, respectively, harbored one or more private clonal driver mutations (**Fig. 1e**) and these values were lower when restricted to untreated metastases (19%, 22% and 22%, respectively). Amongst treated metastases, the proportion of private-clonal drivers increased dramatically across all three cancer types with 71%, 75% and 53% in colorectal, lung and breast cancer, respectively (**Fig. 1e**). This pattern was similarly evident in patients where both untreated and treated metastases were sampled where all (10/10) treated metastases harbored private functional driver mutation(s), but few (2/10) untreated lymph node metastases did (**Table S6**). Therefore, these data suggest that the therapy selects a minor micrometastatic subclone (**Fig. 1g**). In contrast, untreated metastases more commonly originate from the major (or dominant) clone in the primary tumor (**Fig. 1e**). Hence, treatment confers a stringent selective pressure and promotes clonal evolution of the metastasis. Although the overall copy number landscape is highly concordant between paired primary tumors and metastases (**Fig. S10**), copy number analysis revealed a small number of putative driver genes that were more frequently amplified or deleted in metastases relative to primary tumors (increasing from P to M by 15%, **Fig. S11**). These include amplification of *RAC1* and deletions of *FAT1* and *ALB* in colorectal cancer, amplifications of *PLCG1* and *SALL4* and deletions of *NOTCH2, CDKN1B* in lung cancer and amplifications of *IL7R, NIPBL* and deletions of *NOTCH1, PTEN* in breast cancer (**Fig. S11**). Collectively, these data suggest that the genomic drivers required for invasion and metastasis often occur early in the primary tumor (**Fig. 1f**).

### Patterns of metastatic seeding in lymph node and distant metastases

In order to infer the clonality of individual metastases, we compared the CCFs of SSNVs in each P/M pair and the number of M-private clonal SSNVs, P-private clonal SSNVs and P/M shared subclonal SSNVs was denoted as *L*_*m*_, *L*_*p*_ and *W*_*s*_, respectively (**Fig. 2b**). We used the Jaccard similarity index (JSI) where *JSI = w*_*s*_*/*(*L*_*m*_ + *L*_*p*_ + *w*_*s*_) to quantify mutational similarity between P/M pairs ^41^ (**Methods**). Polyclonal seeding is expected to result in a higher JSI than monoclonal seeding due to the higher proportion of shared subclonal SSNVs (higher *W*_*s*_) and the presence of fewer M or P-private clonal SSNVs (lower *L*_*m*_ and *L*_*p*_) (**Fig. 2b**). These patterns were verified by simulation studies using an established agent-based model of spatial tumor progression ^16,30^ (**Figs. S12-S13, Methods**). Notably, polyclonal seeding can be either a multicellular event (by cell cluster) or multiple consecutive monoclonal events (**Fig. 2a**). However, current data is underpowered to distinguish these two scenarios as the resultant patterns of genomic heterogeneity between the primary tumor and metastasis are highly similar. We therefore only modeled polyclonal seeding by cell clusters (**Methods**). By analyzing data from virtual tumors simulated under varied parameters where one biopsy (∼10^6^ cells) was sampled from each primary tumor and metastasis pair (**Methods**), we found that a JSI value of 0.3 maximizes the classification accuracy (91.1%) in distinguishing monoclonal versus polyclonal seeding (**Fig. 2c**). We also simulated MRS data (n=4 samples from each of primary tumor and metastasis) and found that the optimal JSI cutoff increased to 0.4 and yielded an increased classification accuracy (96.3%) relative to single sample data (**Fig. S14**). Given that most (>80%) patients in this study only had only a single sample from the primary tumor and metastasis, we retain the 0.3 cutoff for analyses (**Fig. 2c**). Most metastases exhibited patterns consistent with monoclonal seeding (n=151, 76% of metastases; median JSI=0.075, interquartile range, IQR=0.021–0.138), whereas polyclonal seeding was less frequent (n=48, 24% of metastases; median JSI=0.523, IQR=0.469–0.800) (**Figs. 2c**).

**Figure 2.**
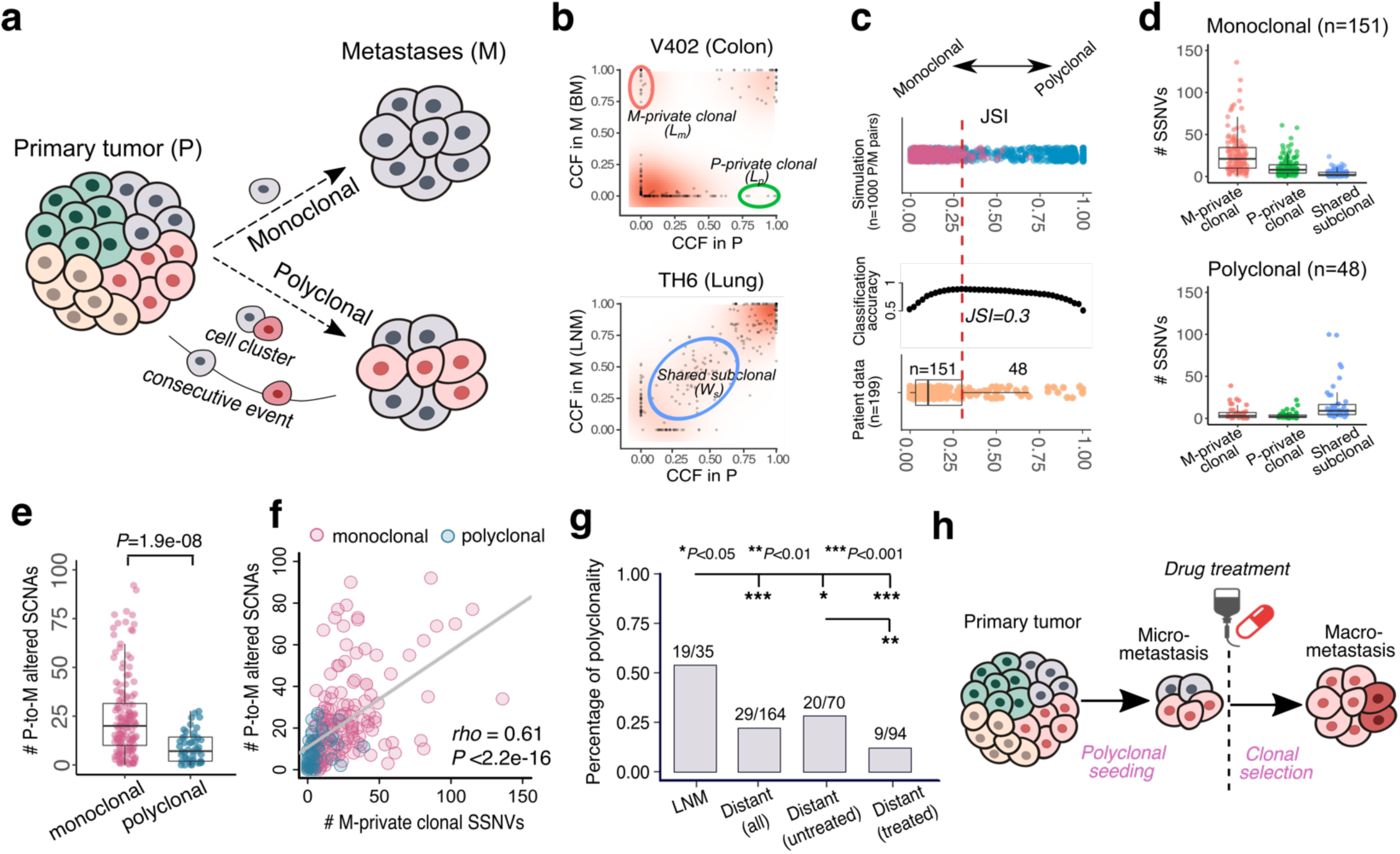
The clonality of lymph node and distant metastases. (**a**) Schematic illustration of monoclonal versus polyclonal seeding for a single metastasis. Polyclonal seeding occurs either through cell cluster or multiple events of single-cell dissemination. (**b**) Distinct patterns of monoclonal versus polyclonal seeding based on the cancer cell fraction (CCF) of SSNVs between P/M pairs. An example patient is shown for each scenario: monoclonal seeding (colon cancer patient V402 with brain metastasis (BM)); polyclonal seeding (lung cancer patient TH6 with lymph node metastasis (LNM)). Green and red circles indicate the P-private clonal SSNVs (the number denoted by *L*_*p*_) and M-private clonal SSNVs (the number denoted by *L*_*m*_), respectively. Blue circle indicates the P/M shared subclonal SSNVs (the number denoted by *W*_*s*_). (**c**) Classification of monoclonal versus polyclonal seeding based on the Jaccard similarity index (JSI). Top, JSI values in 1000 virtual P/M tumor pairs simulated within a spatial tumor growth model with 500 instances of monoclonal seeding (number of metastasis founder cells=1) and 500 instances of polyclonal seeding (number of metastasis founder cells=10). Middle, classification accuracy by varying the cutoff of JSI from 0 to 1 based on the simulation data. Bottom, the JSI values in patient data (n=199 P/M pairs) where the cutoff JSI=0.3 was used to identify monoclonal seeding (n=151) or polyclonal seeding (n=48). (**d**) *L*_*m*_, *L*_*p*_, *W*_*s*_ values in the patient data. Top, monoclonal metastases; bottom, polyclonal metastases. Bar, median; box, 25th to 75th percentile (interquartile range, IQR); vertical line, data within 1.5 times the IQR. (**e**) The number of P-to-M altered SCNAs for monoclonal and polyclonal metastases, respectively. (**f**) Positive correlation between *L*_*m*_ and the number of P-to-M altered SCNAs. n=199 P/M pairs and Spearman’s correlation (*rho*) and *P*-value were reported. (**g**) Polyclonal seeding is common in lymph node metastases (LNM) and untreated distant metastases relative to treated distant metastases. (**h**) Schematic illustration of a scenario where treatment promotes monoclonality as a result of selection for a resistant subclone, despite initial seeded by polyclonal disseminated cells.

As expected, monoclonal metastases (n=151) exhibited significantly higher *L*_*m*_ and *L*_*p*_ values than polyclonal metastases (n=48) (*P*=6.2e-16 and *P*=2.1e-09 for *L*_*m*_ and *L*_*p*_, respectively, two-sided Wilcoxon Rank Sum Test) and significantly lower *W*_*s*_ values (*P*=2.1e-12, two-sided Wilcoxon Rank Sum Test) (**Fig. 2d**). Metastases of monoclonal origin also harbored significantly more SCNAs relative to paired primary tumors than polyclonal metastases (*P*=1.9e-08, two-sided Wilcoxon Rank Sum Test; **Fig. 2e**). Indeed, *L*_*m*_ is highly correlated with the number of P-to-M altered SCNAs (Spearman’s *rho*=0.61, *P*<2.2e-16; **Fig. 2f**), indicating that both SSNVs and SCNAs reflect the clonality of metastases. Polyclonal seeding was more prevalent in axillary lymph node metastases (all, 19/35 or 54%) relative to distant metastases (29/164 or 18%) (*P*=1.8e-05, two-sided Fisher’s exact test; **Figs. 2g and S15**). This pattern is also true for untreated metastases (lymph node vs distant, 17/29 or 59% vs 20/70 or 29%; *P*=0.007, two-sided Fisher’s exact test), potentially reflecting greater lymphatic spread of disseminated cells to the lymph nodes via multiple dissemination events (**Fig. 2a**). Amongst distant metastases, polyclonal seeding was more prevalent in untreated metastases (20/70 or 29%) than treated metastases (9/94 or 10%) (*P*=0.002, two-sided Fisher’s exact test; **Fig. 2g**), presumably because treatment selects for resistant micrometastatic subclones that manifest clinically as monoclonal metastases (**Fig. 2h**). The higher P/M driver gene heterogeneity observed in treated versus untreated metastases (**Fig. 1e**) is consistent with this scenario. The prevalence of polyclonal seeding differed across metastatic sites (lymph node, liver, brain and lung), with brain and lung more commonly exhibiting monoclonal seeding (**Fig. S15b**); these two sites were more commonly biopsied after treatment. In fact, the prevalence of polyclonal seeding amongst lymph node (54%), liver (26%), brain (17%) and lung (8%) metastases is negatively associated with the fraction of metastases that were treated amongst these four sites (17%, 21%, 68%, 92%). This pattern is most evident for brain metastases which had ample numbers of both treated (n=43) and untreated (n=21) metastases. Here, the prevalence of polyclonal seeding is 7% (3/43) amongst treated and 38% (8/21) amongst untreated metastases (*P*=0.004, two-sided Fisher’s exact test; **Fig. S15b**). Therefore, the prevalence of polyclonal seeding is variable across metastatic sites and dependent on whether treatment was administrated before sampling. Since treatment influences the clonality of metastases, we would expect that polyclonal seeding of distant metastases might be more common during the natural course of metastasis (in the absence of treatment) than was observed (18%) here.

We further verified the JSI-based classification of monoclonal versus polyclonal seeding by phylogenetic analysis of patients with MRS of the primary tumor and metastasis (n=13 patients; **Figs. 3** and **S16**). Monoclonal seeding was associated with a monophyletic tree structure (metastatic samples comprise a single phylogenetic clade), whereas polyclonal seeding was associated with a polyphyletic structure (metastatic samples comprise multiple phylogenetic clades) (**Figs. 3** and **S16**).

**Figure 3.**
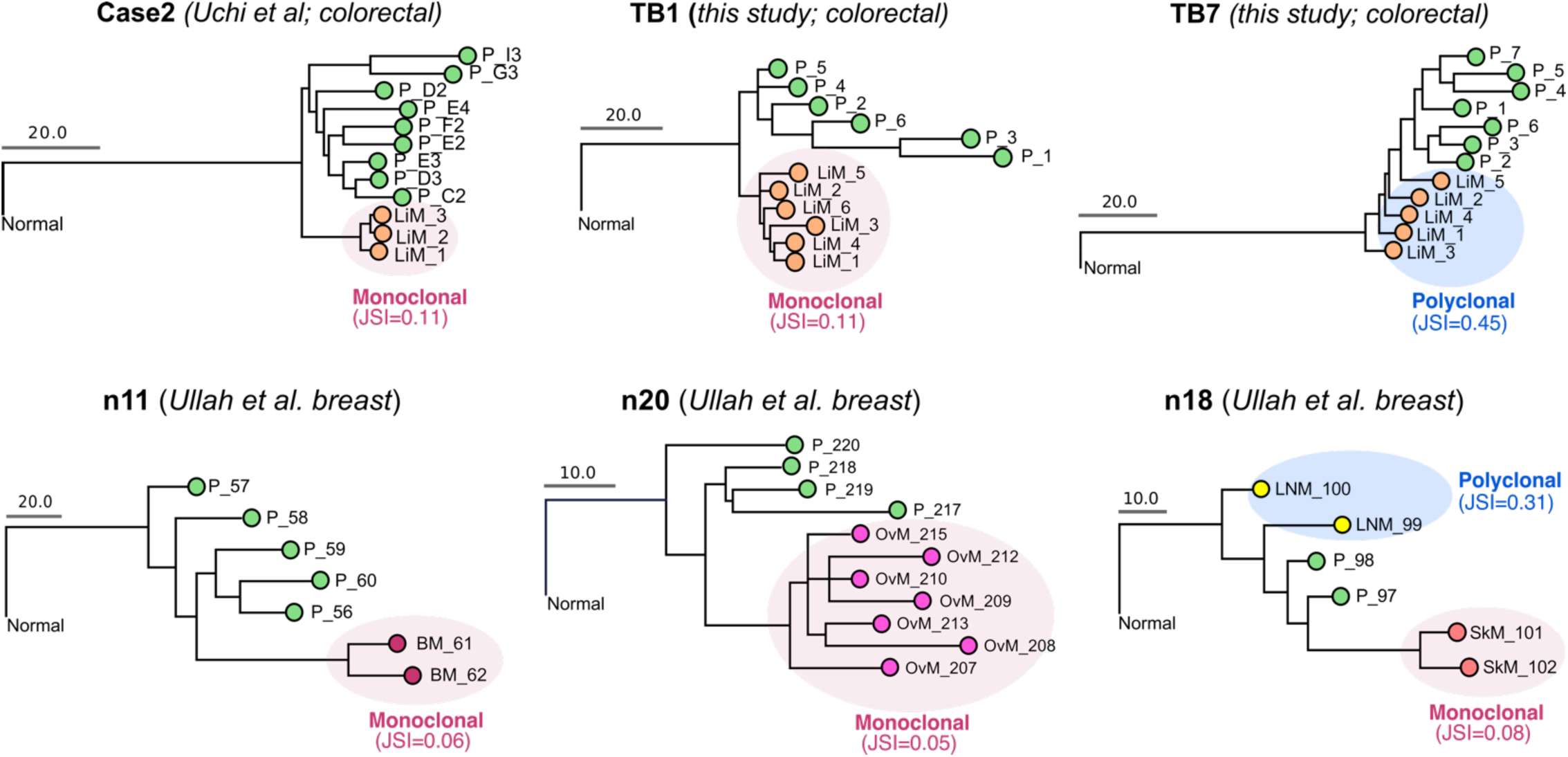
Tumor sample phylogenies based on multi-region sequencing data. The maximum parsimony method was used to reconstruct multi-sample trees for each patient based on the presence or absence SSNVs/indels amongst the samples while accounting for the loss-of-heterozygosity in the mutant sites. For each P/M sample pair, the Jaccard similarity index (JSI) was computed according to Eq. (3) based on the numbers of M-private clonal, P-private clonal and P-M shared subclonal SSNVs. High JSI values (>0.3) indicates polyclonal seeding while low JSI values (≤0.3) indicates monoclonal seeding. Monoclonal seeding gives rise to monophyletic tree structures (pink shading indicates metastatic samples within a single phylogenetic clade), whereas polyclonal seeding gives rise to a polyphyletic structure (blue shading indicates metastatic samples within multiple phylogenetic clades) in the metastasis samples. P, primary tumor; OvM, ovarian metastasis; LNM, lymph node metastasis; SkM, skin metastasis; LiM, liver metastasis. Additional patient data are shown in **Fig. S16**.

We also evaluated the association between clonality and patient outcome (e.g. time to metastatic relapse) based on untreated distant metastases, whereas treated metastases were excluded due the impact of treatment on clonality. In total, there were 70 untreated distant metastases in our cohort: liver: n=45; brain: n=21; bone: n=2; lung: n=1; skin: n=1. We thus focus on liver and brain metastases. Most untreated liver metastases were synchronously diagnosed (91%; 41/45). Of note, all four metachronous liver metastases (time to relapse ranged from 7-8 months) were monoclonal. Amongst brain metastases, 13 exhibited patterns consistent with monoclonal seeding while 8 were consistent with polyclonal seeding. Amongst, 31% (4/13) and 37.5% (3/8) were metachronous, respectively. Notably, the time to relapse was longer for monoclonal brain metastasis (median=26 months, IQR=(19, 36)) than polyclonal brain metastasis (median=11 months, IQR=(9, 17)). Although limited by the small sample size, this (non-significant) trend suggests that polyclonal seeding may be associated with worse prognosis. However, further studies on large (untreated) metastatic cohorts are warranted.

### Chronology of metastatic seeding

Previously, we described a computational framework (SCIMET) to estimate the timing of metastatic seeding relative to primary tumor size based on MRS of P/M pairs ^16^. Application of this approach to colorectal cancer yielded quantitative evidence for early systemic spread, well before the primary tumor was clinically detectable. Since MRS data was not available for the vast majority of patients in this cohort, we developed a new computational method that leverages exome sequencing data from a single biopsy to time metastatic seeding (**Figs. 4a** and **S17, Supplementary Note**). The time (in years) from metastatic seeding to diagnosis of the primary tumor (*t*_*s*_) can be approximated by:

**Figure 4.**
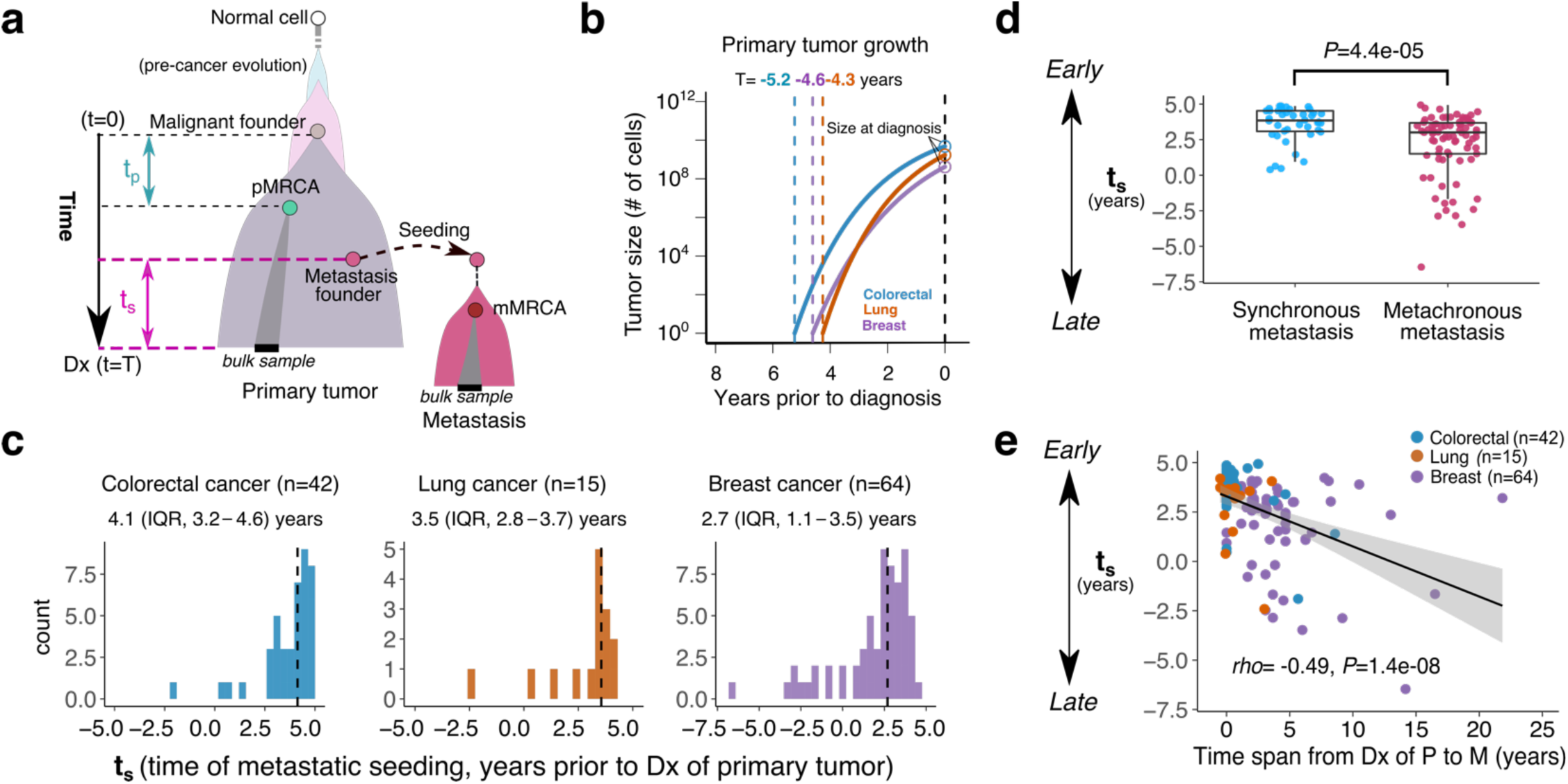
Chronology of metastatic seeding. (**a**) Schematic for the timing of metastatic seeding prior to diagnosis of the primary tumor in number of years, *t*_*s*_. *T* denotes the total time of primary tumor expansion from emergence of the malignant founder cell to diagnosis while *t*_*p*_ denotes the time from emergence of the malignant founder cell to the most recent common ancestor (MRCA) of cells in primary bulk sample (denoted pMRCA). *t*_*s*_ can be estimated by *Eq.(1)*. Dx, diagnosis (**b**) Estimation of the average *T* with a Gompertzian growth model is 5.2 (interquartile range or IQR, 4.3–7.7), 4.3 (IQR, 2.7–4.4) and 4.6 (IQR, 3.2–6.6) years for colorectal, lung and breast cancer, respectively. (**c**) Estimation of the time of metastatic seeding (*t*_*s*_) for individual distant metastases (monoclonal) in each cancer types. The median *t*_*s*_ and IQR are shown. Negative *t*_*s*_ indicates that the metastasis was seeded after the diagnosis of primary tumor. (**d**) The distribution of *t*_*s*_ in synchronous metastases (n=41) and metachronous metastases (n=80). *P*-value, Wilcoxon Rank-Sum Test (two-sided). Bar, median; box, 25th to 75th percentile (IQR); vertical line, data within 1.5 times the IQR. (**e**) Correlation between *t*_*s*_ and the time span from diagnosis of primary tumor to metastasis. Spearman’s correlation (rho) and *P*-value are reported.

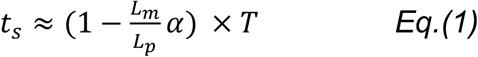

where *L*_*m*_ and *L*_*p*_ correspond to the number of M-private clonal SSNVs and P-private clonal SSNVs, respectively; *T* is the primary tumor expansion age (time from emergence of carcinoma founder cell to diagnosis); α *=* t_*p*_*/T* where *t*_*p*_ is the time from emergence of carcinoma founder cell to the most recent common ancestor in the primary tumor sample (pMRCA, **Figs. 4a** and **S17, Supplementary Note**). The time fraction α is expected to be small because bulk sequencing only detects relatively high frequency mutations that occur early during tumor growth or that are strongly selected for ^42-44^. We applied our established agent-based model of spatial tumor growth ^30^ to simulate a large set of virtual tumors (n=1000, each ∼10^9^ cells) with varying growth rates (**Methods**). *In silico* sequencing of a single biopsy (each ∼10^6^ cells, mean depth=100X) from the virtual tumors (n=1000) yields an estimate of 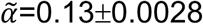 (**Fig. S17**), confirming the observation that bulk sequencing typically only detects high-frequency mutations that occur early during tumor growth. Here we assume a model of stringent selection (selection coefficient, s=0.1) during growth of the primary tumor based on our prior analysis of MRS data which revealed evidence for selection in primary colon cancers within a metastatic cohort ^16^, as well as in primary lung ^30^ and breast ^45^ cancers. This assumption is further supported by the finding that most primary tumors in this cohort (57/65 or 88% evaluable tumors) exhibited variant allelic frequencies (VAF) distributions that were not consistent with neutral evolution ^46^, despite limitations of this analysis (**Fig. S18; Methods**).

We utilized a Gompertzian model of tumor growth ^47^, to estimate the tumor expansion age (*T*) for each of the three cancer types (**Supplementary Note**) where tumor size and doubling time (DT) at diagnosis were obtained from literature review (**Table S8**). This yields estimates of average tumor expansion age of 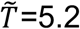 (IQR, 4.3–7.7), 4.3 (IQR, 2.7–4.4) and 4.6 (IQR, 3.2–6.6) years for colorectal, lung and breast cancer, respectively (**Fig. 4b** and **Table S9**). Chronological estimates of seeding time relative to diagnosis of the primary tumor 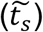 can be computed by *Eq*.(1) as follows: 4.1 years (IQR, 3.2–4.6), 3.6 years (IQR, 2.8–3.7) and 2.7 years (IQR, 1.1–3.5) for colorectal, lung and breast cancers, respectively (**Fig. 4c** and **Table S9**). The estimated timing of metastasis here 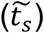 agreed with our previous estimates (using the colorectal cancer cohort) of primary tumor size at time of metastatic seeding ^16^ (Spearman’s *rho*= −0.55, *P*=0.014, **Fig. S19;** note the negative correlation since this we estimate backward time and the previous study estimated forward time). Of note, while [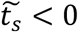 may indicate metastatic seeding after diagnosis/resection of the primary tumor, large *L*_*m*_ values can lead to 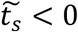 see *Eq.*(1)) even when the metastasis was seeded before diagnosis of the primary tumor. To mitigate this uncertainty, samples with estimated seeding times later than the actual time of diagnosis of metastasis were excluded (n=12 for breast, 1 for colorectal and 1 for lung cancer, respectively) (**Supplementary Note**). We find that 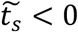 was more common in breast cancer and more generally breast cancers disseminated closer to the time of detection (later) compared to colorectal and lung cancers (**Fig. 4c**). This may be because screening mammography detects relatively small primary breast tumors (<2 cm) ^48^.However, even after normalization to primary tumor age (namely *t*_*s*_/*T*), which depends on tumor size and the underlying growth parameters (**Supplementary Note**), breast cancer was found to disseminate later than colorectal and lung cancers (**Fig. S20**). Most breast cancer metastases (83%) in this cohort were biopsied after adjuvant therapy (**Fig. S1**), whereas this fraction is fewer in colorectal (13%) and lung (20%) cancer metastases and breast cancers harbored more private driver mutations than colorectal and lung cancers (**Figs. 1a-c**). Thus, the genomic complexity of metastatic relapses in breast cancer relative to unpaired early-stage primary tumors ^12^ at least in part reflects the selective effect of treatment on the genome, rather than the drivers of metastatic spread. Of note, HER2-positive breast cancers tended to disseminate earlier than HER2-negative breast cancers (**Fig. S21**) consistent with this subgroup having the highest risk of distant metastasis before the routine use of adjuvant trastuzumab ^33^, which has revolutionized the treatment of this disease in part by targeting occult micrometastases.

As expected, metachronous metastases were often seeded later than synchronous metastases (median *t*_*s*_=3.8 vs 3.0, *P*=5.6e-05, two-sided Wilcoxon Rank-Sum Test; **Fig. 4d**). In fact, *t*_*s*_ was highly correlated with the clinical time span from diagnosis of primary tumor to metastasis (**Fig. 4e**), indicating that metastases that manifest late clinically were seeded later. Since primary tumor size at diagnosis is an important predictor of a patient’s prognosis (time to metastatic relapse) (**Fig. S22a**), we suspect that metastases in patients with larger primary tumor size at initial diagnosis were seeded earlier (namely larger *t*_*s*_). Indeed, *t*_*s*_ is positively associated with the primary tumor size at diagnosis (Spearman’s *rho*=0.24, *P*=0.023; **Fig. S22b**). These results corroborate our estimates of metastatic timing. According to *Eq.*(1), a larger number of M-private clonal mutations (larger *L*_*m*_) indicates later dissemination. Supporting this theory, metachronous metastases showed significantly larger *L*_*m*_ than synchronous metastases (metachronous: median *L*_*m*_=24, IQR=16–40; synchronous: median *L*_*m*_=11, IQR=6–32; *P*=6.5e-4, two-sided Wilcoxon Rank-Sum Test; **Fig. S23a**). This pattern held for SCNAs where metachronous metastases showed significantly more SCNAs relative to the primary tumor as compared to synchronous metastases (**Fig. S23b**). Since metachronous metastases were generally seeded later than synchronous metastases (**Fig. 4d**), this is consistent with the higher degree of genomic divergence with primary tumor in late seeded metastases ^18^. Given that adjuvant treatment targets micrometastases, presumably delaying clinical metastasis, the timing of metastatic seeding of metachronous metastases following treatment might be even earlier. Collectively, these data indicate that systemic spread can occur several years prior to diagnosis of the primary tumor but with variability across histologies and subgroups.

## Discussion

We performed a systematic analysis of exome sequencing data in paired primary tumors and metastases across three common cancers: colorectal, lung and breast and find that polyclonal seeding is common in lymph node metastases (19/35, 54%; most untreated) and untreated distant metastases (20/70, 29%), but rare (9/94, 10%) in metastases sampled after adjuvant therapy (**Fig. 2g**). Consistent with these results, treated metastases were strongly enriched for functional driver mutations as compared to untreated metastases (**Fig. 1e**). This finding indicates that driver gene heterogeneity is minimal between untreated metastases and primary tumors (**Fig. 1e**). Comparisons of paired primary tumors and distant metastases indicates that systemic spread can occur rapidly following malignant transformation, often several years prior to diagnosis of the primary tumor across three major types (**Fig. 4c**). These results are consistent with other reports of early seeding based on animal models and disseminated tumor cells ^9,49,50^.

Our analyses on driver gene heterogeneity, clonality and the timing of metastases provide important insights into the clonal dynamics of metastatic progression. First, in the absence of treatment, metastases often arise from the major clone in the primary tumor and lack metastasis-specific driver mutations (**Fig. 1f**). Consistent with these observations, a recent multi-cancer study demonstrated that driver gene heterogeneity is also minimal amongst multiple untreated metastases within individual patients ^51^. Moreover, the prevalence of polyclonal seeding in untreated lymph node and distant metastases indicates multiple cell subpopulations in primary tumor have acquired the metastatic competence. Half of all metastases (51%) studied here were biopsied after treatment, and these commonly exhibited monoclonal seeding accompanied by private driver mutations. As such, polyclonal seeding may be relatively common, but the ultimate pattern of clonality in the metastatic lesion is influenced by treatment.

Second, our quantitative framework demonstrates that systemic spread typically begins 2-4 years prior to the diagnosis of primary tumor (**Fig. 4c**). These data suggest that in some patients, metastatic seeding can happen very early especially for synchronously diagnosed metastases (**Figs. 4e, 5a**). Metachronous distant metastases following treatment occurred relatively later than synchronous distant metastases and harbored more genomic aberrations and driver mutations (**Figs. 1e and S23**). These data imply that treatment remodels the clonal evolution of metastasis and may select disseminated cells harboring drug resistant mutations (**Fig. 5a-b**). As such metastasis-specific mutations are unlikely to be the drivers of metastasis, but instead are associated with resistance (**Fig. 5b**). This interpretation is of clinical relevance and helps to clarify the observation that metastatic relapses are more genomically complex than unpaired early-stage primary breast tumors ^12^. At the same time, adjuvant therapy directed at micrometastatic disease is effective for many patients, at least for a period of time, thus forestalling disease progression ^52^. Unfortunately, in cases where relapse occurs, the resultant metastatic outgrowth may be driven by a more aggressive, treatment resistant clone.

**Figure 5.**
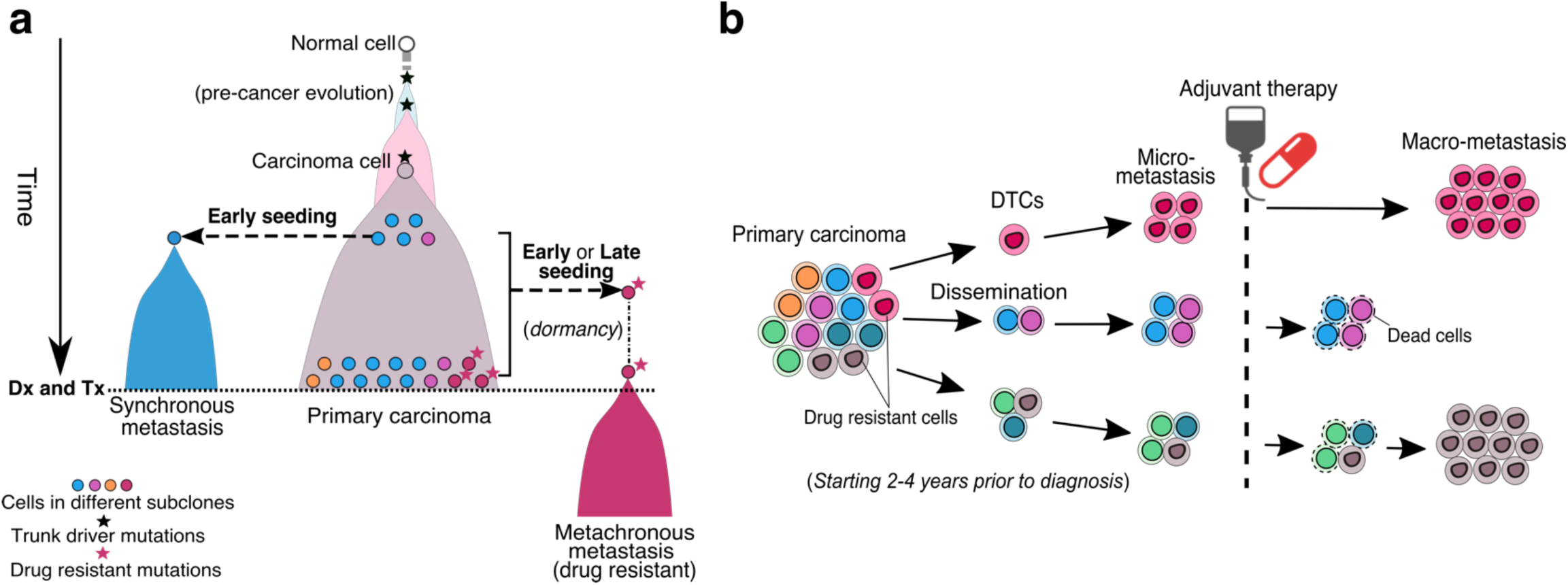
Schematic model of metastatic spread and the impact of therapy. **(a)** Schematic illustration of early versus late metastatic seeding leading to synchronous and metachronous metastases. Metastatic seeding starts quickly following the emergence of the founding carcinoma cell. Synchronous metastasis, which exhibits low genomic divergence with primary tumor, is seeded early by the major clone in primary tumor. Metachronous metastasis, exhibit higher genomic divergence relative to the primary tumor and often emerge after adjuvant therapy. Metachronous metastases with specific driver mutations that confer resistance can be selected leading to high genomic divergence between the primary tumor and treated metastasis. Dx, diagnosis; Tx, treatment. (**b**) Treatment (here adjuvant therapy) remodels the clonal architecture of metastasis. Dissemination and metastatic seeding (monoclonal or polyclonal) initially gives rise to undetectable micrometastases. While treatment may eliminate drug-sensitive micrometastatic lesions, those that are resistant grow out. Metastatic relapse following adjuvant treatment may be delayed by adjuvant treatment, but this may result in a more aggressive, resistant lesion. DTCs, disseminated tumor cells.

To our knowledge, this study is based on the largest collection of paired primary tumors and metastases across multiple cancer types with genomic data, but several limitations remain. First, the majority of tumors (>80% for P or M) were sequenced to standard depth (median=88, IQR=(65, 110), **Table S3**), which is likely underpowered to identify polyclonal seeding patterns based on shared subclonal (low frequency) mutations (**Fig. 2b**). Simulations show that multi-region sequencing (n=4 from each of P and M) increases the accuracy of classifying monoclonal and polyclonal seeding as compared to single sample. Second, more than half of distant metastases were biopsied after drug treatment which can substantially remodel the clonal architecture of the metastasis by promoting monoclonality and genomic divergence. If these two main confounders are considered, we would expect that polyclonal seeding of distant metastases is more common than inferred (18%) here and that metastatic dissemination might occur even earlier. Our findings highlight the importance of studying the natural course of metastasis as well as the impact of therapy on this process. Future studies of paired primary tumors and metastases with comprehensive treatment information subject to dense multi-region sampling and single cell sequencing may provide additional resolution on these processes.

## Acknowledgments

We thank Hang Xu, Katherine McNamara, Eran Kotler, Jennifer Caswell-Jin and other members of Curtis laboratory for valuable discussions. We thank Jiguang Wang and Quanhua Mu for providing the scripts for the ternary plot. C.C. is supported by the National Institutes of Health through the NIH Director’s Pioneer Award (DP1-CA238296), the American Association for Cancer Research (AACR) and the Emerson Collective. Z.H is supported by an Innovative Genomics Initiative (IGI) Postdoctoral Fellowship.

## Author contributions

Z.H and C.C conceived and designed the study. Z.H performed all computational analyses. Z.L reviewed the published studies, extracted and analyzed the clinical data. Z.M processed the in-house clinical samples and generated the genomic data. Z.H and C.C wrote the manuscript, which was reviewed by all authors.

## Methods

### Whole-exome sequencing (WES) of paired primary tumors and metastases

We performed a comprehensive review on the published studies through surveying the PubMed database (https://www.ncbi.nlm.nih.gov/pubmed/), in which whole-exome sequencing (WES) was performed for matched normal tissues, primary tumors (P) and metastases (M) in the same patients. We focused on colorectal, lung and breast cancers given the availability of large patient data in these three cancer types. In total, the raw sequencing reads data for 586 tumor samples from 181 patients in 13 published studies were accessed and retrieved (**Table S1**). We also generated multi-region sequencing (MRS) data for two colorectal cancer patients (mCRCTB1 and mCRCTB7) with liver metastases from whom excess de-identified tissue was collected during the course of routine care, hence this is not considered human subjects research. A total of n=5-7 regions were sequenced for these two P/M pairs resulting in 24 tumor samples. Here tumor tissues with cellularity >60% were selected for DNA isolation using the QIAamp DNA FFPE Tissue Kit (Qiagen) and libraries were generated using the Agilent SureSelect Human All Exon kit for sequencing on the Illumina Hiseq 2500. Clinical information was retrieved from the original studies, including patient age at initial diagnosis, time span from initial diagnosis of primary tumor to diagnosis of metastasis, treated information and subtype (**Table S2-S3**). We define synchronous metastases if the time span between diagnosis of primary tumor and metastasis is within 3 months and metachronous metastases if the time span is ≥3 months.

An established bioinformatics pipeline was used to detect somatic single nucleotide variations (SSNVs), small insertions/deletions (indels) and somatic copy number alterations (SCNAs), estimate tumor purity/ploidy and estimate the cancer cell fraction (CCF) for each SSNVs/indels in corresponding samples ^16,30^. In particular, paired sequencing reads were aligned to human reference genome (NCBI build hg19) with BWA (v.0.7.10) ^53^. Duplicate reads were marked with Picard Tools (v.1.111). Aligned reads were further processed with GATK 3.4.0 for local re-alignment around insertions and deletions and base quality recalibration.

### SSNVs and indel calling

SSNVs were called by MuTect (v.1.1.7) ^54^ for each tumor/normal pair. SSNVs failing MuTect’s internal filters, having fewer than 10 total reads or 3 variant reads in the tumor sample, fewer than 10 total reads in the normal sample, or mapping to paralogous genomic regions were removed. Additional Varscan (v.2.3.9) ^55^ filters were applied to remove SSNVs with low average variant base qualities, low average mapping qualities among variant supporting reads, strand bias among variant supporting reads and high average mismatch base quality sums among variant supporting reads, either within each tumor sample or across all tumor samples from the same patient. The maximal observed variant allele frequencies (VAF) across all samples from each patient were calculated based on raw output files from MuTect. SSNVs with maximal observed VAFs lower than 0.05 were removed. For FFPE specimens, additional filters were applied to exclude possible artifactual SSNVs. Specifically, artifacts among C>T/G>A SSNVs with bias in read pair orientation were filtered in each individual FFPE sample, similar to the approach of Costello *et al* ^56^. Of note, 26%, 80% and 81% of primary colorectal, lung and breast cancer samples were FFPE while 29%, 50% and 55% of metastatic samples in these three cancer types were FFPE. FFPE artifacts are at low frequency in primary tumors (median VAF= 0.056–0.085 across studies) and metastases (median VAF= 0.017–0.090 across studies). On average, more than 70% of the FFPE artifacts across the cohort were specific to the primary tumors, consistent the primary tumor more commonly being FFPE than the metastasis. We also sought to exploit the multi-sample information in the same patients to retrieve read counts for SSNVs. To obtain the depth and VAF information across all samples from the same patient, for each SSNV and in each tumor sample that an SSNV was not originally called in, the total reads and variant supporting reads were counted using the *mpileup* command in SAMtools (v.1.2) ^57^. Only reads with mapping quality ≥ 40 and base quality at the SSNV locus ≥ 20 were counted and used to calculate the VAF for that SSNV. Small insertions/deletions (indels) were called with Strelka (v.1.0.14) ^58^. SSNVs and indels were annotated with ANNOVAR (v.20150617) ^59^ and those in protein coding regions were retained for downstream analyses.

### Copy number analysis

Copy number analysis was performed using TitanCNA (v.1.5.7) ^60^. Briefly, TitanCNA uses depth ratio and B-allele frequency information to estimate allele-specific absolute copy numbers with a hidden Markov model, and estimates tumor purity and clonal frequencies. Only autosomes were used in copy number analysis. First, for each patient, germline heterozygous SNP at dbSNP 138 loci were identified using SAMtools and SnpEff (v.3.6) in the normal sample. HMMcopy (v.0.99.0) ^61^ was used to generate read counts for 1000bp bins across the genome for all tumor samples. TitanCNA was used to calculate allelic ratios at the germline heterozygous SNP loci in the tumor sample and depth ratios between the tumor sample and the normal sample in bins containing those SNP loci. Only SNP loci within WES covered regions were then used to estimate allele-specific absolute copy number profiles. TitanCNA was run with different numbers of subclones (n=1-3). One run was chosen for each tumor sample based on visual inspection of fitted results, with preference given to the results with a single subclone unless results with multiple subclones had visibly better fit to the data. Results from tumor samples from the same patient were inspected together to ensure consistency. Overall ploidy and purity for each tumor sample was calculated from the TitanCNA results.

Differentially altered SCNAs in the metastasis relative to paired primary tumor (P-to-M) were identified if following three criteria were satisfied simultaneously: 1) absolute copy number in the metastasis was larger than 2.8 or less than 1.2; 2) copy number relative to median ploidy in the metastasis was larger than 0.8 or less than −0.8; 3) changes relative to the primary tumor in both absolute copy number and relative copy number were larger than 0.8 or less than −0.8. For multi-region sequencing data, segmented log depth ratios (adjusted for purity and ploidy) for each primary CRC and paired metastasis were averaged across multiple-regions from the same tumor site.

### Cancer cell fraction (CCF) estimates and identification of clonal and subclonal mutations

The CCFs and their variation (95% confidence interval or 95% CI) for each SSNVs/indels in the corresponding samples were estimated with CHAT (v 1.0) ^62^. CHAT includes a function to estimate the CCF of each SSNVs by adjusting its variant allele frequency (VAF) based on local allele-specific copy numbers at the SSNV locus. SSNV frequencies and copy number profiles estimated from previous steps were used to calculate the CCFs for all SSNVs in autosomes. The CCFs were also adjusted for tumor purity using the estimates by TitanCNA. In brief, for an SSNV residing in a genomic segment with a total copy number of *CN*_*t*_, minor allele copy number of *CN*_*b*_ and cellular prevalence *P*_*CNA*_of the CNA in the tumor content, the estimated *CCF* of the SSNV is:

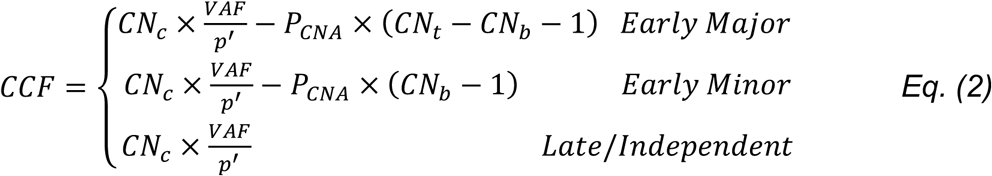

where *CN*_*C*_ *= CN*_*t*_ × *P*_*CNA*_ + 2 × (1 − *P*_*CNA*_) and the effective purity 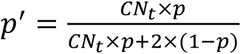 (*p* is estimated tumor purity) and VAF is the observed variant allele frequency. The temporal ordering and background composition of SSNVs and SCNAs was inferred by comparing the conditional probabilities of the observed number of mutant reads out of total reads, under each scenario and CNA configuration (*CN*_*t*_, *CN*_*b*_, *P*_*CNA*_) as follows: *Early Major* or *Mino*r: SSNV in the major or minor allele occurred before the CNA; *Late*: SSNV occurred after the CNA; *Independent*: the SSNV and CNA occurred in independent lineages ^62^.

To distinguish clonal and subclonal SSNVs/indels in each sample, we employ the following criterion: clonal – 95% CI overlaps with 1; subclonal – the upper bound of 95% CI is smaller than 1, as previously used ^63^.

Since bulk sequencing data is underpowered to detect low frequency mutations, determining whether a mutation is truly private mutations to one site is challenging. Thus, “private” SSNV/indels in one site relative to another site is paratactically defined as the CCF<5% in another site as our previous study ^16^. For multi-region sequencing data, the merged CCFs by integrating multiple regions were used:

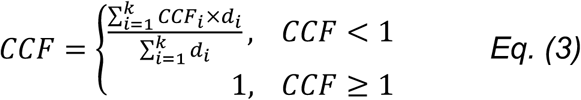

where *d*_*i*_ and *CCF*_*i*_ are the sequencing depth and CCF estimation in region *i*, respectively.

### Sample quality control for downstream analysis

The CCFs of SSNV/indels for each P/M sample pair were visualized using the scatter plot and manually checked in order to identify problematic samples. In particular, for each P/M pair, a cluster of SSNV/indels centered around CCF=1 is expected which represent truncal (P/M shared clonal) mutations that occurred prior to malignant transformation of the founding cell in the primary tumor. The patients (n=5) with none of or very few (<10) trunk SSNVs/indels were excluded as which implies independent (non-clonal) origin for the primary tumor and metastasis. Furthermore, patients (n=42) with a diffusely distributed cluster for truncal SSNVs/indels were also excluded since this is likely caused by low tumor purity or low sequencing quality. After these filtering steps, 457 tumor samples from 136 metastatic cancer patients including 39 colorectal cancers (181 tumor samples), 30 lung cancer (74 tumor samples) and 67 breast cancers (202 tumor samples) were retained for downstream analysis in this study. Regarding the histological subtypes, all colorectal cancers were microsatellite stable (MSS). For lung cancer, 67% (20/30) were adenocarcinoma, 30% (9/30) were squamous carcinoma, while 3% (1/30) were small cell lung cancer. For breast cancer, 6% (4/67) were ER+/HER2+, 6% (4/67) were ER-/HER2+, 51% (34/67) were ER+/HER2-, 19% (13/67) were triple negative (TN), while 18% (12/67) were unknown (**Table S2**). Of note, there is a bias towards obtaining more paired primary and distant metastases from triple negative (TN) breast tumors since they tend to recur earlier than ER+ tumors (many within 5 years), where for some subsets of ER+/HER2-disease there is a persistent risk of recurrence up to two decades after diagnosis ^33,64^.

### Jaccard similarity index

The number of M-private clonal, P-private clonal and P-M shared subclonal SSNVs for each P/M pair was denoted as *L*_*m*_, *L*_*p*_ and *W*_*s*_ respectively. For two sets, the Jaccard similarity index (JSI) is defined for the intersection divided by the union of these two sets. Thus, the JSI for a P/M pair can be defined as:

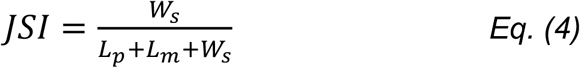

For multi-region sequencing data, *L*_*m*_, *L*_*p*_ and *W*_*s*_ was counted by pairwise comparison of each sample pair from the P and M. The mean *L*_*m*_, *L*_*p*_ and *W*_*s*_ was used to compute the JSI by *Eq.(4).*

### Functional assessment of non-silent somatic mutations

To identify functional driver gene mutations, three commonly used computational methods, PolyPhen-2 ^36^ (http://genetics.bwh.harvard.edu/pph2/), FATHMM-XF ^37^ (http://fathmm.biocompute.org.uk/fathmm-xf/)and CHASMplus^38^ (https://karchinlab.github.io/CHASMplus/), were utilized to perform the function (“driverness”) assessment on the nonsynonymous SSNVs amongst putative cancer genes derived from TCGA pan-cancer ^65^ and COSMIC (Release v87, Nov. 13, 2018). Stopgain/splicing point mutations and indels on putative cancer genes are classified as functional drivers automatically.

Putative cancer genes were curated by merging all TCGA pan-cancer drivers (n=299) ^65^ and additional cancer type-specific drivers annotated by COSMIC Cancer Gene Census (https://cancer.sanger.ac.uk/cosmic; n=47, 40 and 9 for colorectal, lung and breast cancers, respectively). For PolyPhen-2, a SSNV is considered as “functional” when the functional report (“pph2_class”) is “deleterious”. For FATHMM-XF, a SSNV is considered as “functional” when the functional report (“Warning”) is “pathogenic”. For CHASMplus, a SSNV is considered as “functional” when the FDR < 0.05. In this study, the SSNVs, predicted to be functional by any of these three methods, were considered as functional mutations. Metascape ^66^ (http://metascape.org) was used to perform gene ontology (GO) analysis of functional driver genes.

### Driver enrichment analysis

Clonal non-silent SSNVs/indels in a metastatic lesion can be considered truncal clonal (or P-M shared clonal) or M-private clonal where the number is denoted *L*_*s*_*_total* and *L*_*m_*_*total*, respectively. Meanwhile, the functional driver SSNVs/indels in a metastasis are denoted *L*_*s*_*_driver* and *L*_*m*_*_driver*, respectively. The ratios, *L*_*s*_*_total/L*_*m_*_*total and L*_*s*_*_driver*/*L*_*m*_*_driver*, can be evaluated for functional enrichment of drivers on the truncal or M-private branch of the corresponding phylogenetic tree. Since *L*_*s*_*_driver* and *L*_*m*_*_driver* are small values (*L*_*m*_*_driver ∼* 0 for many metastases), they lead to high variation in the *L*_*s*_*_driver*/*L*_*m*_*_driver* ratio. A down-sampling (bootstrapping) step (50% of the patients each time) was performed in which sampled patient data were merged to derive the *L*_*s*_*_total/L*_*m_*_*total and L*_*s*_*_driver*/*L*_*m*_*_driver* ratios. 100 repeated down-samplings were performed for each of the three cancer types to derive statistical measures.

### Mutational signatures, dN/dS and test of neutrality

MuSiCa ^67^ (http://bioinfo.ciberehd.org:3838/MuSiCa/) was used to extract mutation signatures based on non-negative matrix factorization ^68^ for P/M shared clonal (truncal) SSNVs, M-private clonal SSNVs and M-private subclonal SSNVs respectively, in each of the three cancer types. dndscv ^40^ (https://github.com/im3sanger/dndscv) was used to compute the ratio of nonsynonymous and synonymous SSNVs (dN/dS) for missense and nonsense mutations, respectively and for P/M shared clonal (trunk) SSNVs, M-private clonal SSNVs and M-private subclonal SSNVs, respectively, in each of the three cancer types. We evaluated whether a tumor exhibits a pattern consistent with neutral evolution or strong selection during growth by analyzing the variant frequency distribution (VAF) of subclonal SSNVs. Under neutral evolution, the number of subclonal SSNVs with VAF larger than *f* in a tumor cell population follows a power-law distribution: *m*(*f*)∼1/*f* ^46^. The adjusted VAFs (equivalent to CCF/2) for subclonal SSNVs (in the range of 0.1−0.3) were used here and only tumors with at least 20 subclonal SSNVs in this range were analyzed (n=65 primary tumors and 79 metastases). By fitting this model and using a threshold of R^2^=0.98, the mode of evolution (neutral or selection) can be inferred (**Fig. S18**). There are notable limitations to this analysis, including the lack of MRS data and the fact that many primary tumors were FFPE. Nonetheless, the finding that the majority of patients exhibit primary tumor VAF distributions consistent with subclonal selection, is in-line with our prior reports in a metastatic colorectal cancer cohort with MRS data, where the majority of primary colon cancers exhibited evidence of subclonal selection, consistent with the metastatic clone having a selective growth advantage ^16^. Additionally, analysis of multi-region sequencing data suggest that subclonal selection may be relatively common in primary lung ^30^ and breast ^45^ cancers.

### Phylogenetic tree reconstruction

We ran PHYLIP ^69^ via an online version http://www.trex.uqam.ca/index.php?action=phylip&apP=dnapars)andappliedthe Maximum Parsimony method to reconstruct the phylogeny of multiple specimens from individual patients based on the presence or absence of SSNVs/indels. The SSNVs/indels residing a region with different loss-of-heterozygosity (LOH) status between paired primary tumor and metastasis were filtered, since which may lead to erroneous presence or absence of SSNVs/indels in paired P and M. When multiple maximum parsimony trees werereported,wechosethetoprankedsolution.FigTree (http://tree.bio.ed.ac.uk/software/Figuretree/) was employed to visualize the reconstructed trees.

### Spatial agent-based modeling of metastatic progression

We employed our previously established three-dimensional agent-based tumor evolution framework ^30^ to model tumor growth, mutation accumulation and metastatic dissemination after malignant transformation. Pre-malignant clonal expansions prior to transformation do not contribute to the genetic heterogeneity of an established tumor (since all such alterations are clonal), and thus were not modeled since dissemination is assumed to occur after malignant transformation of the founding carcinoma cell. In this model, spatial tumor growth is simulated via the expansion of deme subpopulations (composed of ∼5k cells with diploid genome), mimicking the glandular structures often found in epithelial tumors and metastases and consistent with the number of cells found in individual colorectal cancer glands (∼2,000-10,000 cells). The deme subpopulations expand within a defined 3D cubic lattice (Moore neighborhood, 26 neighbors), via peripheral growth while cells within each deme are well-mixed without spatial constraints and grow via a random birth-and-death process (division probability *b* and death probability *d*=1-*b* at each generation). Once a deme exceeds the maximum size (10,000 cells), it splits into two offspring demes via random sampling of cells from a binomial distribution (*Nc*, 0.5), where *Nc* is the current deme size.

To model monoclonal seeding, a single cell at the tumor periphery was randomly sampled as the metastasis founder cell. To model polyclonal seeding, a cluster of cells (n=10) were randomly sampled from the whole tumor in order to maximize the clonal diversity within the metastasis founder cells. This is because if the clonal diversity in the metastasis founder cells is low, it essentially models the scenario of monoclonal seeding by a cluster of genetically similar cells. The metastasis grows at same spatial model with primary tumor started from the metastasis founder cell or cell cluster (n=10). During each cell division in the growth of primary tumor and metastasis, the number of neutral passenger mutations acquired in the coding portion of the genome follows a Poisson distribution with mean *u*. Thus, the probability that *k* mutations occurred in each cell division is as follows:

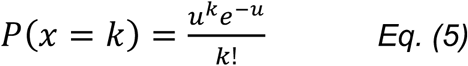

where an infinite sites model and constant mutation rate are assumed during tumor progression. Advantageous mutations also arise stochastically via a Poisson process with mean *u*_*s*_ during each cell division. We assume *u*_*s*_=10^−5^ per cell division in the genome and each increases the cell division probability ^70^. The cell birth and death probabilities for a selectively beneficial clone are *b*_*s*_=*b*×(1+*s*) and *d*_*s*_=1-*d*_*s*_=1-*b*×(1+*s*), respectively, thus the selective advantage for an advantageous mutation is defined as *s*=*b*_*s*_/*b*-1.

During simulation of primary and metastatic growth, each mutation is assigned a unique index that is recorded with respect to its genealogy and host cells, enabling analysis of the mutational frequency in a bulk sample of tumor cells during different stages of growth. We simulate growth until the primary and metastasis reach a size of ∼10^9^ cells (or ∼10 cm^3^) and then sample a bulk subpopulation (consisting of ∼10^6^ cells) at the peripheral region of the primary tumor and metastasis, respectively. The VAF of all SSNVs in the sampled bulk subpopulation is considered the true VAF (denoted by *f*_*T*_), whereas the observed allele frequency is obtained via a statistical model that mimics the random sampling of alleles during sequencing. Specifically, we employ a Binomial distribution (*n, f*_*T*_) to generate the observed VAF at each site given its true frequency *f*_*T*_ and number of covered reads *n*. The number of covered reads at each site is assumed to follow a negative-binomial distribution (*Negative Binomial*(*size, depth*)) where depth is the mean sequencing depth and size corresponds to the variation parameter. We assume *depth*=100 and *size*=2 for the sequencing data in each tumor region and tissue purity=0.6 in order to model normal cell contamination in clinical samples. A mutation is called when the number of variant reads is ≥3, thereby applying the same criteria as for the patient tumors.

We employed a mutation rate *u*=0.6 per cell division in the exonic region (corresponding to 10^−8^ per site per cell division in the 60Mb diploid coding regions). In order to model varying scenarios of tumor growth dynamics, selection and timing of metastatic dissemination, for each primary tumor/metastasis (P/M) pair, the birth probability *b* of founding cells, selection coefficient *s* and primary tumor size at dissemination *N*_*d*_ was sampled from a uniform distribution, *b*∼U(0.55, 0.65), log10(*s*)∼U(−3,-1) and log10(*N*_*d*_)∼U(4,8), respectively. 500 virtual P/M pairs were simulated under each of the monoclonal seeding and polyclonal seeding scenarios. The number of M-private clonal SSNVs (*L*_*m*_), P-private clonal SSNVs (*L*_*p*_) and P/M shared subclonal SSNVs (*W*_*s*_) for each P/M pair were counted from the simulation data and the simulated JSI was computed by *Eq.(4).*

## Data availability

The exome sequencing data for in-house collected colorectal cancer patients have been deposited at the European Genotype Phenotype Archive (EGA) under accession number EGAS00001003573. The accession numbers for public datasets are listed in Table S1.

## Code availability

Code used for genomic data analysis are available from: https://github.com/cancersysbio

