## Supplemental Information for "Multi-cancer analysis of clonality and the timing of systemic spread in paired primary tumors and metastases"

*Supplementary Figures 1-23 (page 2-24)*
*Supplementary Note (page 25-27)*
*Supplementary Tables 8-9 (page 28-29)*
*Supplementary References (page 30-31)*

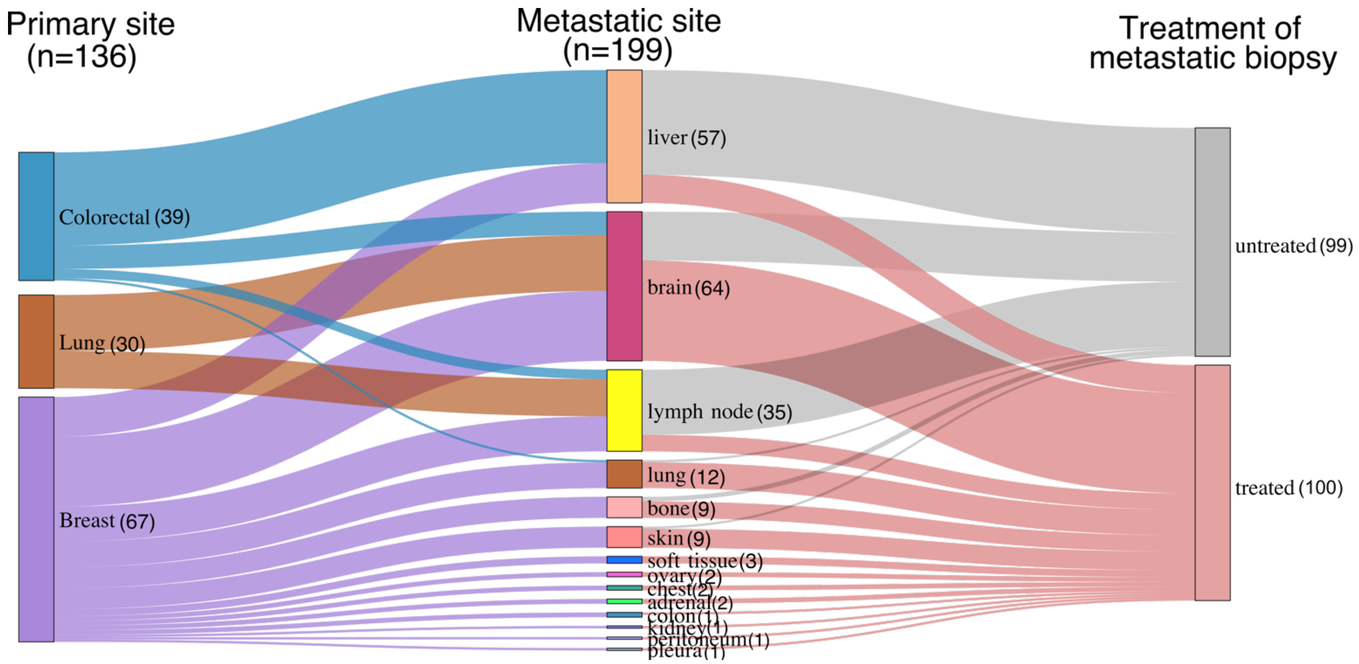

**Supplementary Figure 1. Sankey diagram of patient cohorts with paired primary tumors and metastases.** In total, 136 primary tumors and 199 matched metastases from colorectal, lung and breast cancers were included. Treatment status is indicated.

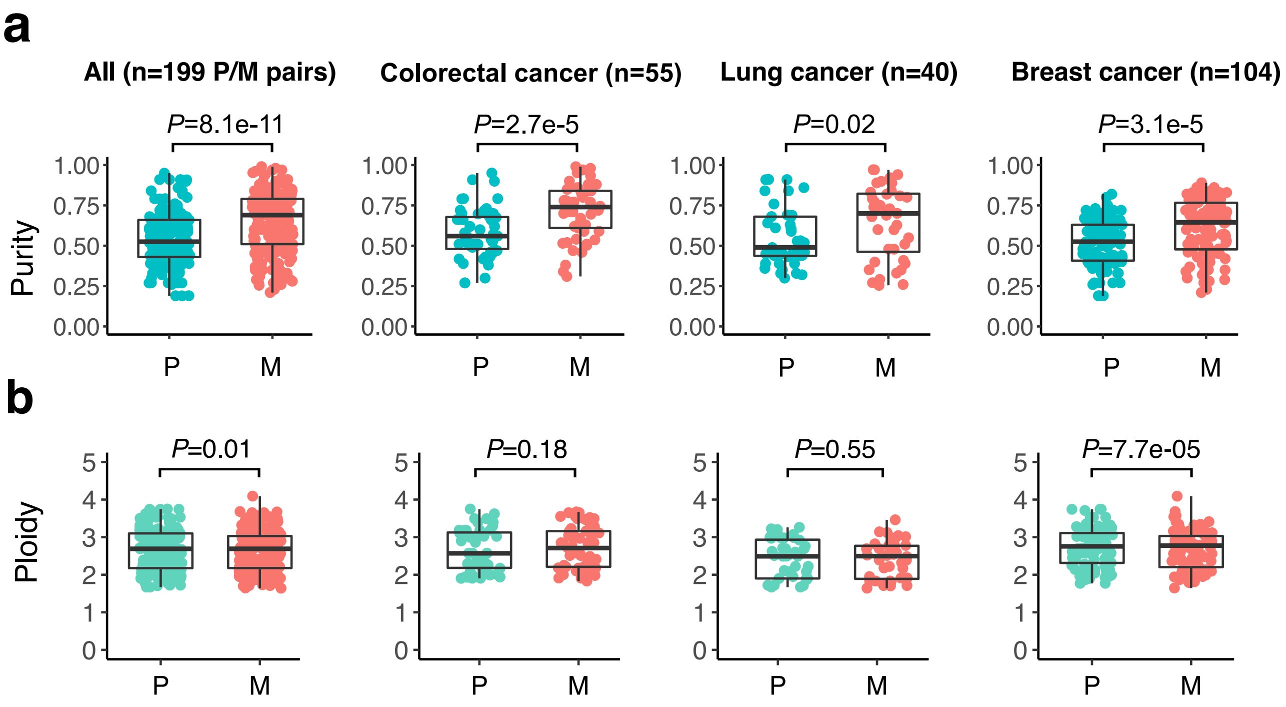

**Supplementary Figure 2. Estimated purity and ploidy in paired primary tumors and metastases.** *P*-value, Wilcoxon Rank-Sum Test (two-sided, paired data). Bar, median; box, 25th to 75th percentile (interquartile range, IQR); vertical line, data within 1.5 times the IQR. Mean purity and ploidy across samples were used for cases with multi-region sequencing data.

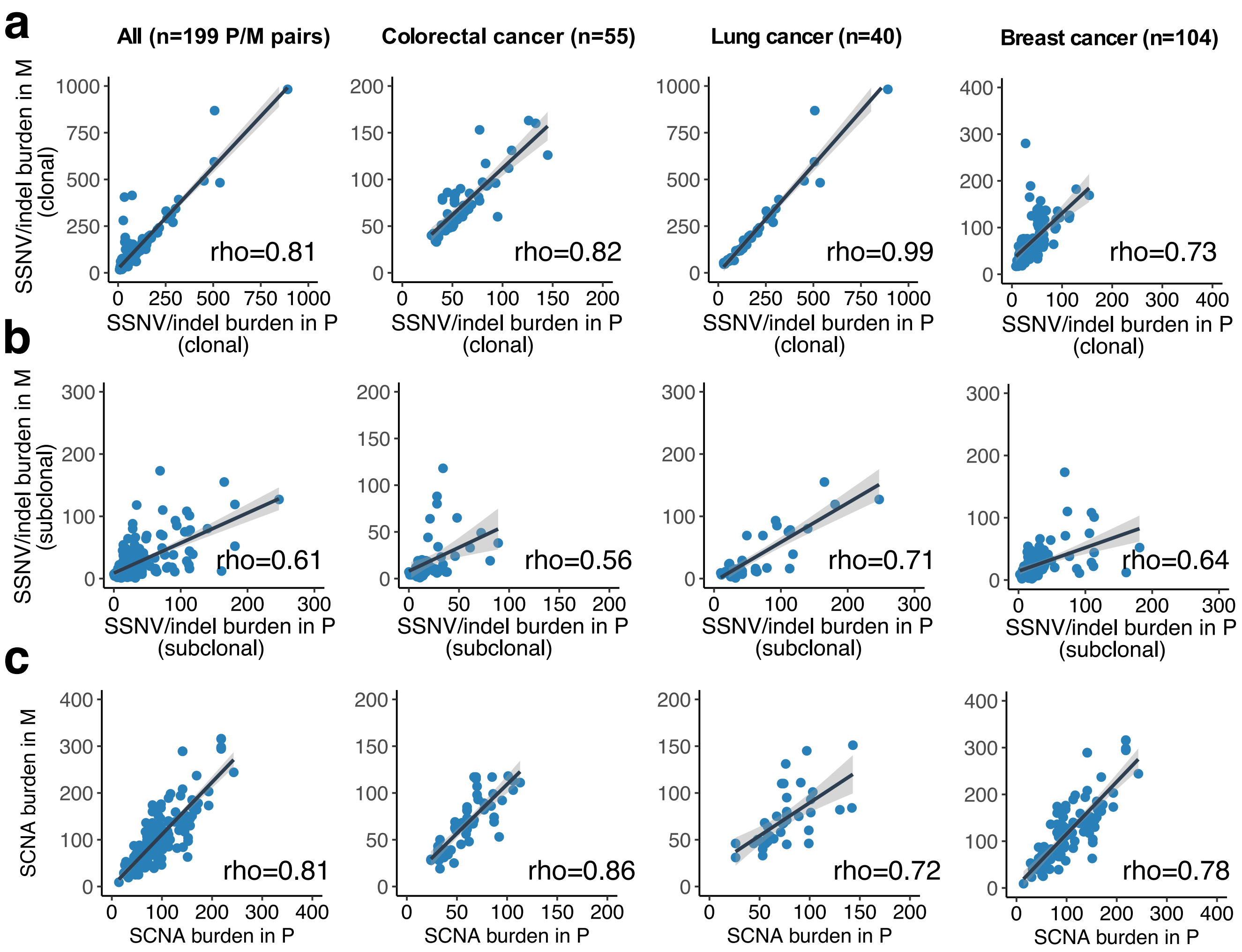

**Supplementary Figure 3. Concordance of mutation burden in paired primary tumors (P) and metastasis (M).** Concordance amongst (**a**) *Clonal* SSNVs (**b**) *Subclonal* SSNVs and (**c**) SCNAs are indicated. Spearman's correlation (*rho)* is reported. Mean mutation burden across samples was used for multi-region sequencing data.

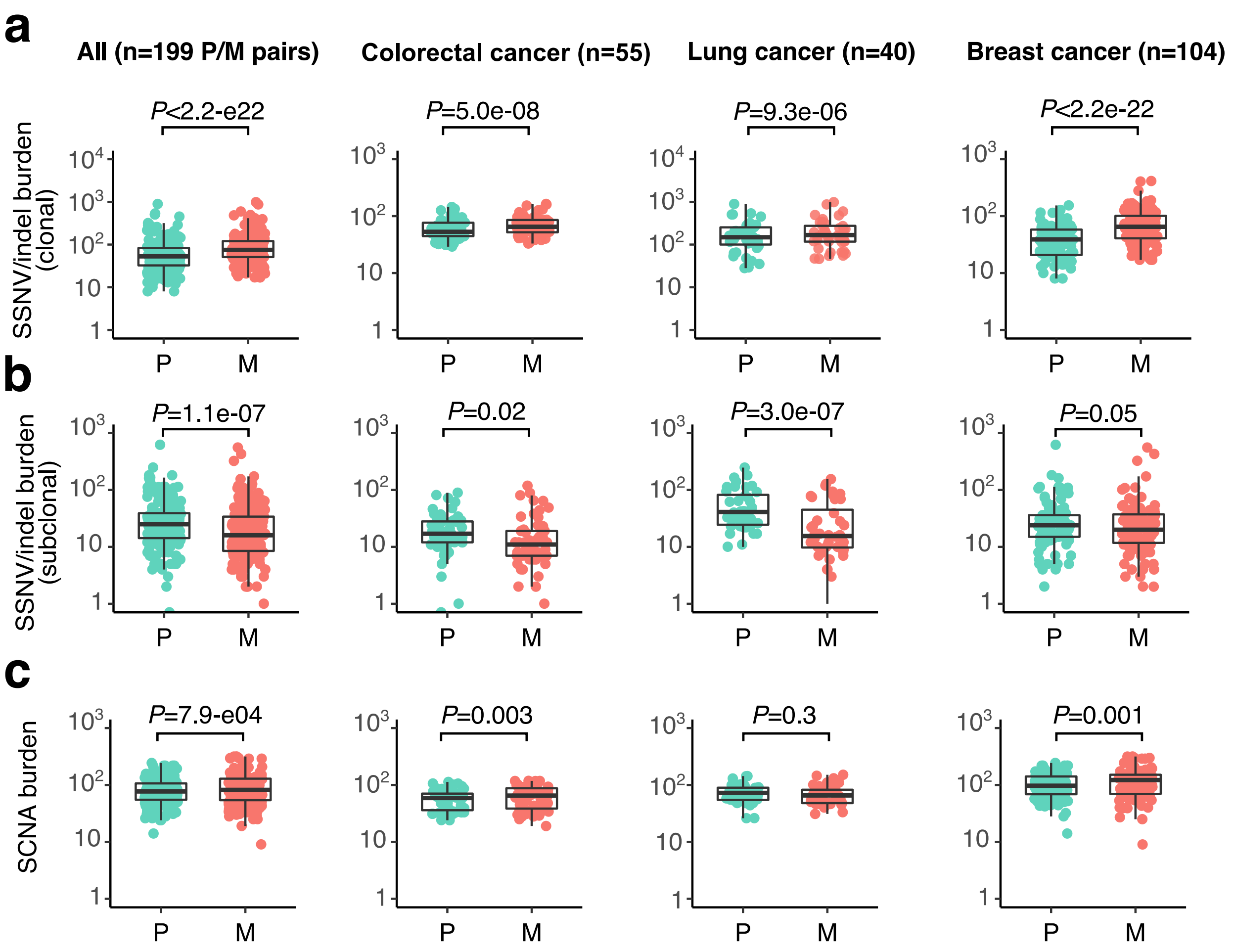

**Supplementary Figure 4. Metastases exhibited slightly higher clonal SSNV and SCNA burden but lower subclonal SSNV burden than primary tumors.** Concordance amongst (**a**) *Clonal* SSNVs (**b**) *Subclonal* SSNVs and (**c**) SCNAs. *P*-value, Wilcoxon Rank-Sum Test (two-sided, paired data). Bar, median; box, 25th to 75th percentile (interquartile range, IQR); vertical line, data within 1.5 times the IQR. Mean mutation burden across samples was used for multi-region sequencing data.

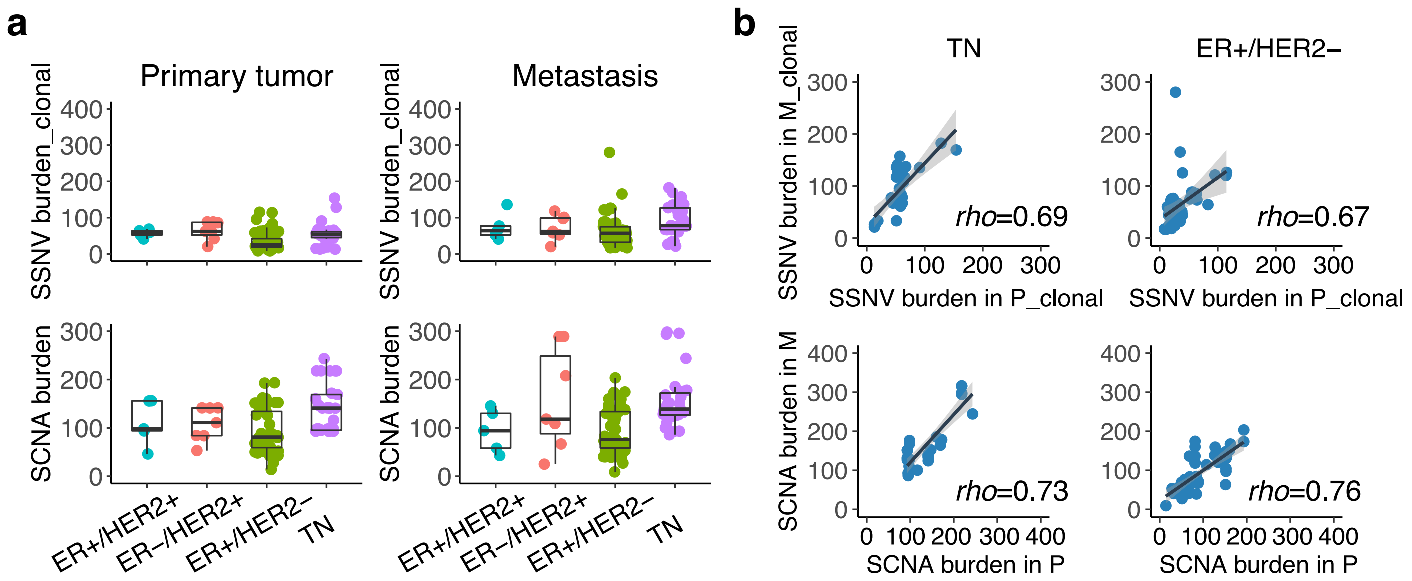

**Supplementary Figure 5. Mutational burden in paired primary breast tumors and metastases by histopathologic subtype**. (**a**) The burden of SSNVs (clonal) or SCNAs in paired primary tumors (P) and metastases (M) in the four histopathologic breast cancer subtypes (number of P/M pairs, ER+/HER2+: n=5; ER- /HER2+: n=7; ER+/HER2-: n=48; TN: n=29). (b) Correlation between the burden of SSNVs (clonal) and SCNAs in paired primary tumors and metastases in TN and ER+/HER2- subtypes, respectively. Spearman's correlation (*rho)* is reported.

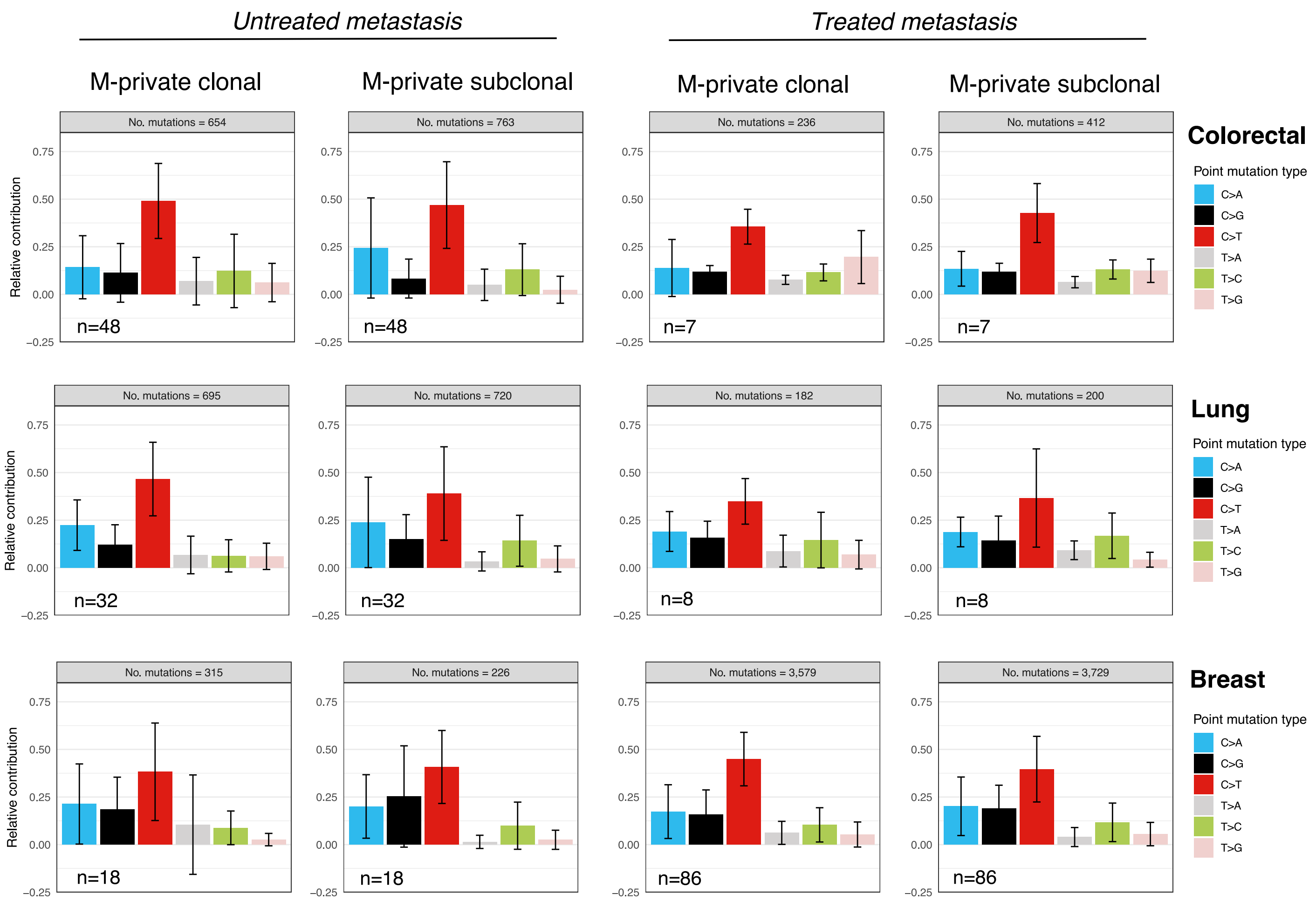

**Supplementary Figure 6. Mutational spectrum for M-private SSNVs.** The relative contribution of six classes of substitutions amongst all mutations in treated or untreated metastases and for M-private clonal or subclonal SSNVs across colon, lung and breast cancers.

M-private *clonal* driver mutations (n=132)

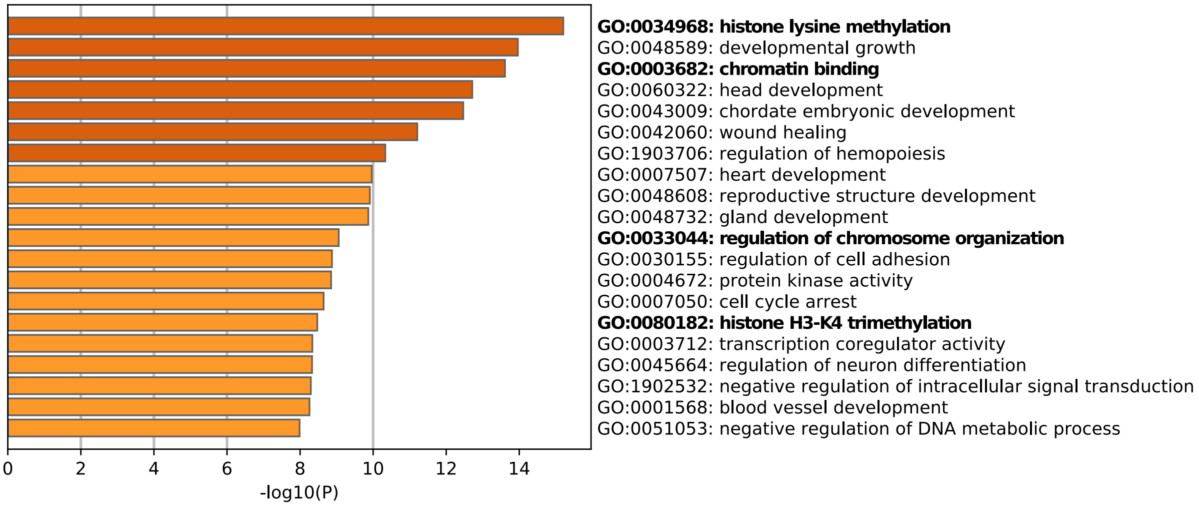

M-private *subclonal* driver mutations (n=131)

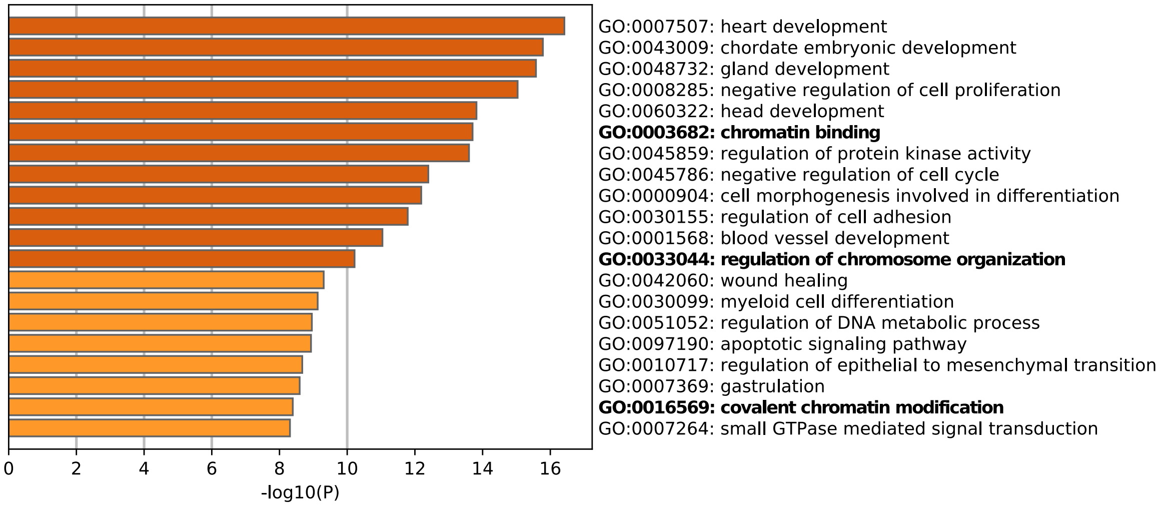

**Supplementary Figure 7. Gene ontology (GO) enrichment for M-private driver mutations.** Functional driver mutations that were private to the metastatic lesion across the three cancer types were pooled for GO enrichment analysis (top panel: clonal drivers; bottom panel: subclonal drivers). The top 20 most significant GO terms are shown; chromatin-related terms denoted in bold. The full list of significant terms and corresponding genes are shown in **Table S7**.

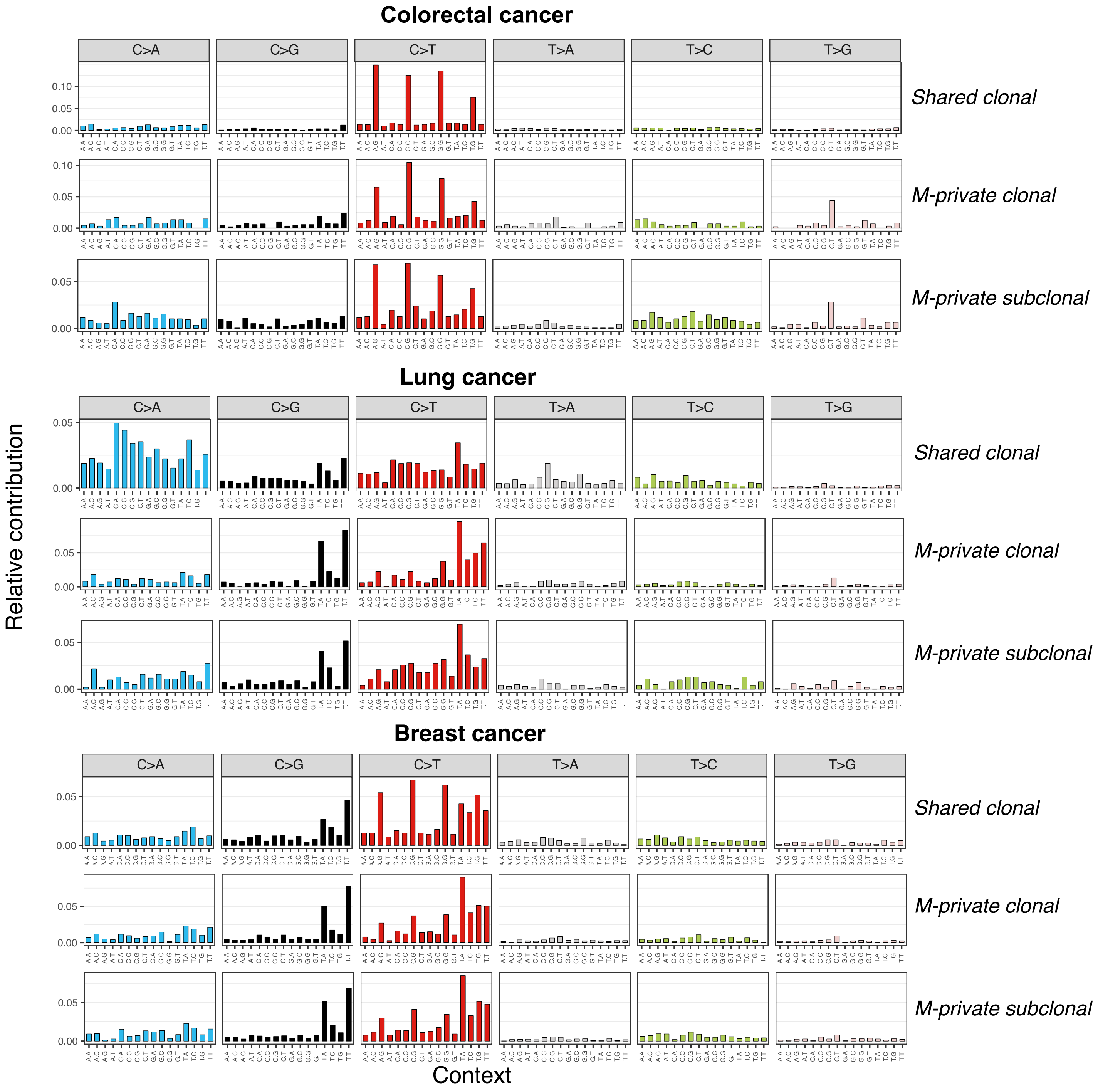

**Supplementary Figure 8. The mutation spectrum of SSNVs.** The bars indicate the relative contribution for each of the 96 context-dependent substitutions for P/M shared clonal, M-private clonal and M-private subclonal SSNVs, respectively.

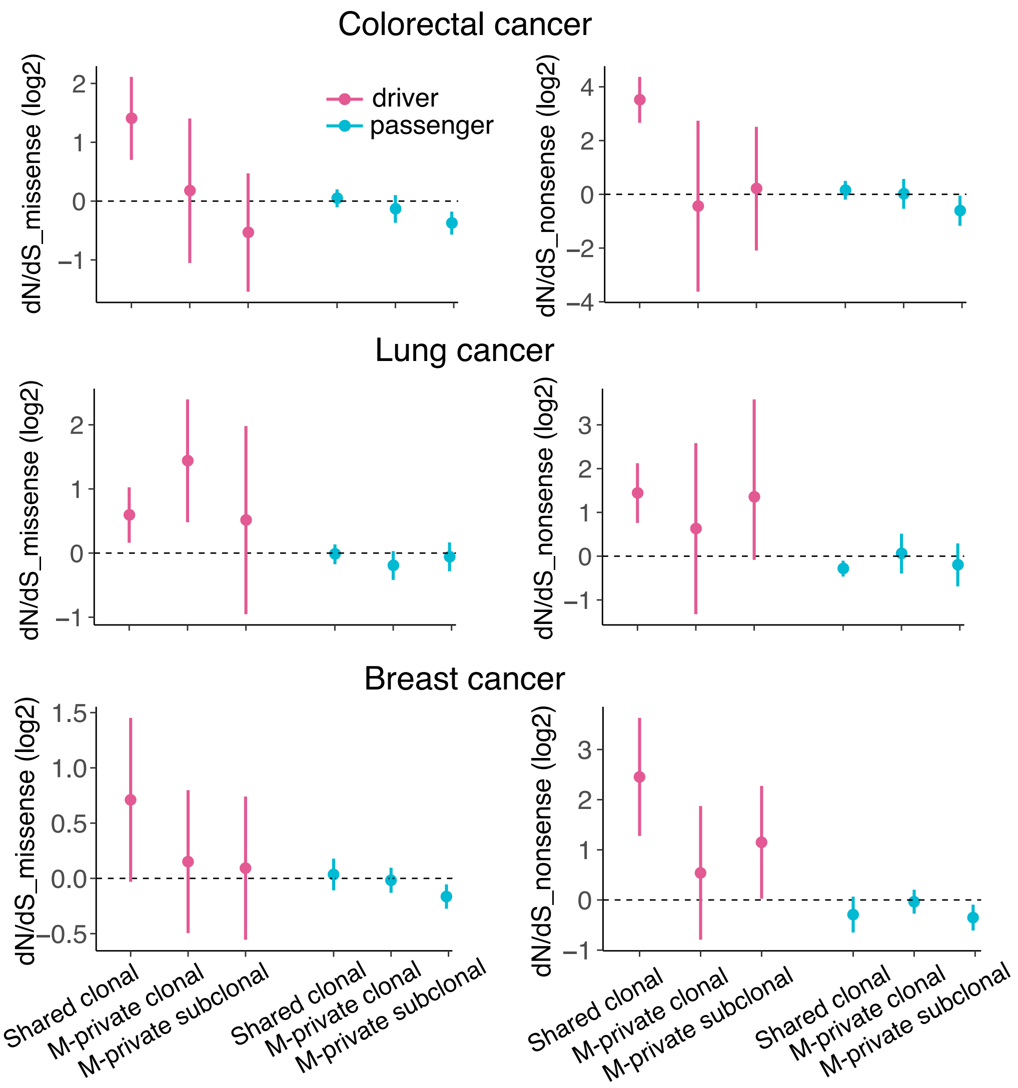

**Supplementary Figure 9. The ratio of nonsynonymous to synonymous mutations, dN/dS.** The dN/dS ratios of missense mutations (left panel) or nonsense mutations (right panel) relative to synonymous mutations are shown (in the scale of log2). The dN/dS ratios for putative driver genes and passengers were computed separately. The driver gene list was obtained by merging TCGA pan-cancer drivers and COSMIC Cancer Gene Census (**Methods**). Circles and vertical lines correspond to the mean and 95% CI of the dN/dS ratio, respectively.

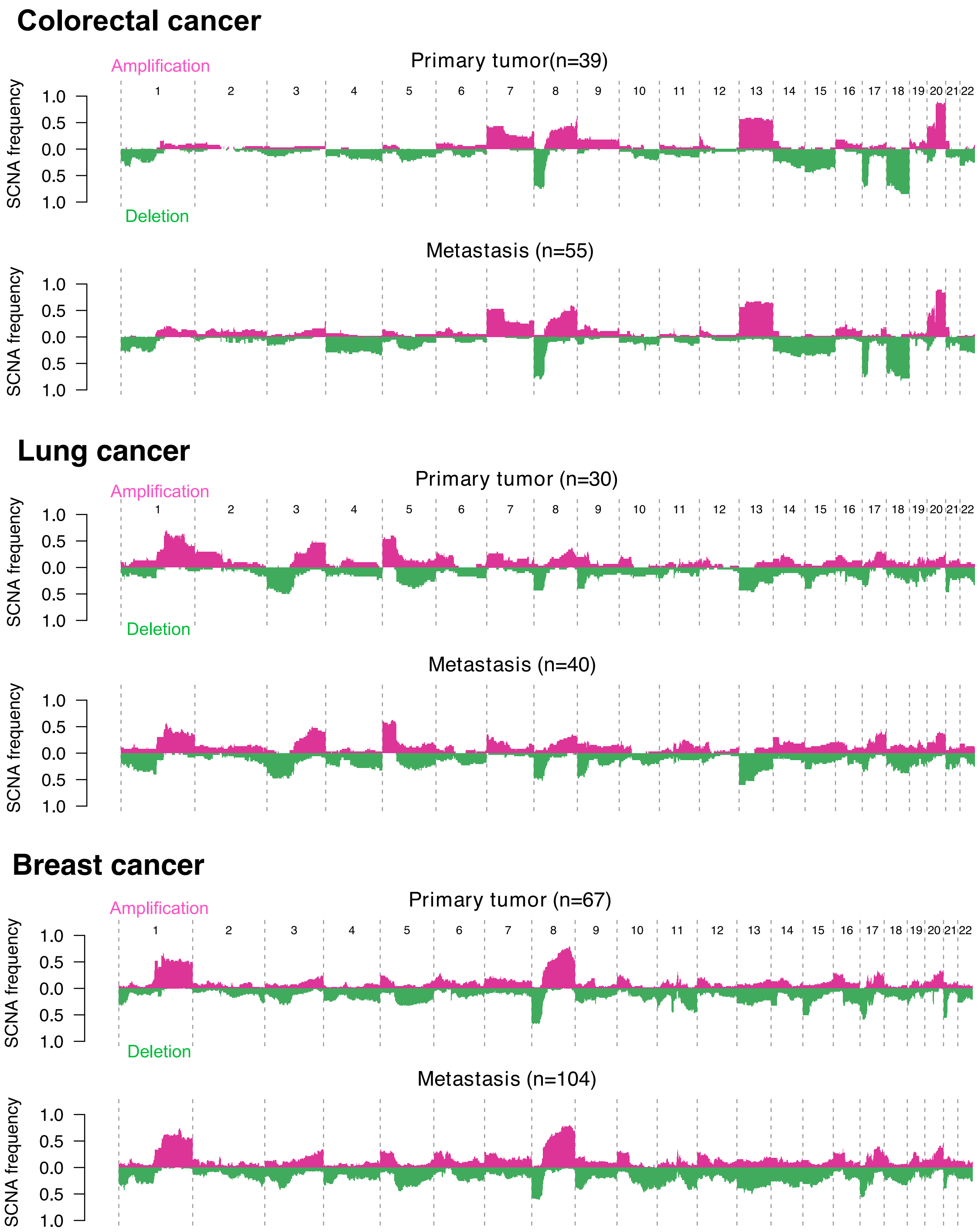

**Supplementary Figure 10. The SCNA frequency for primary tumors and metastasis across three cancer types.** The frequency of somatic copy number amplification or deletion for each genomic bin (1 Mb) is shown for primary tumors and metastases.

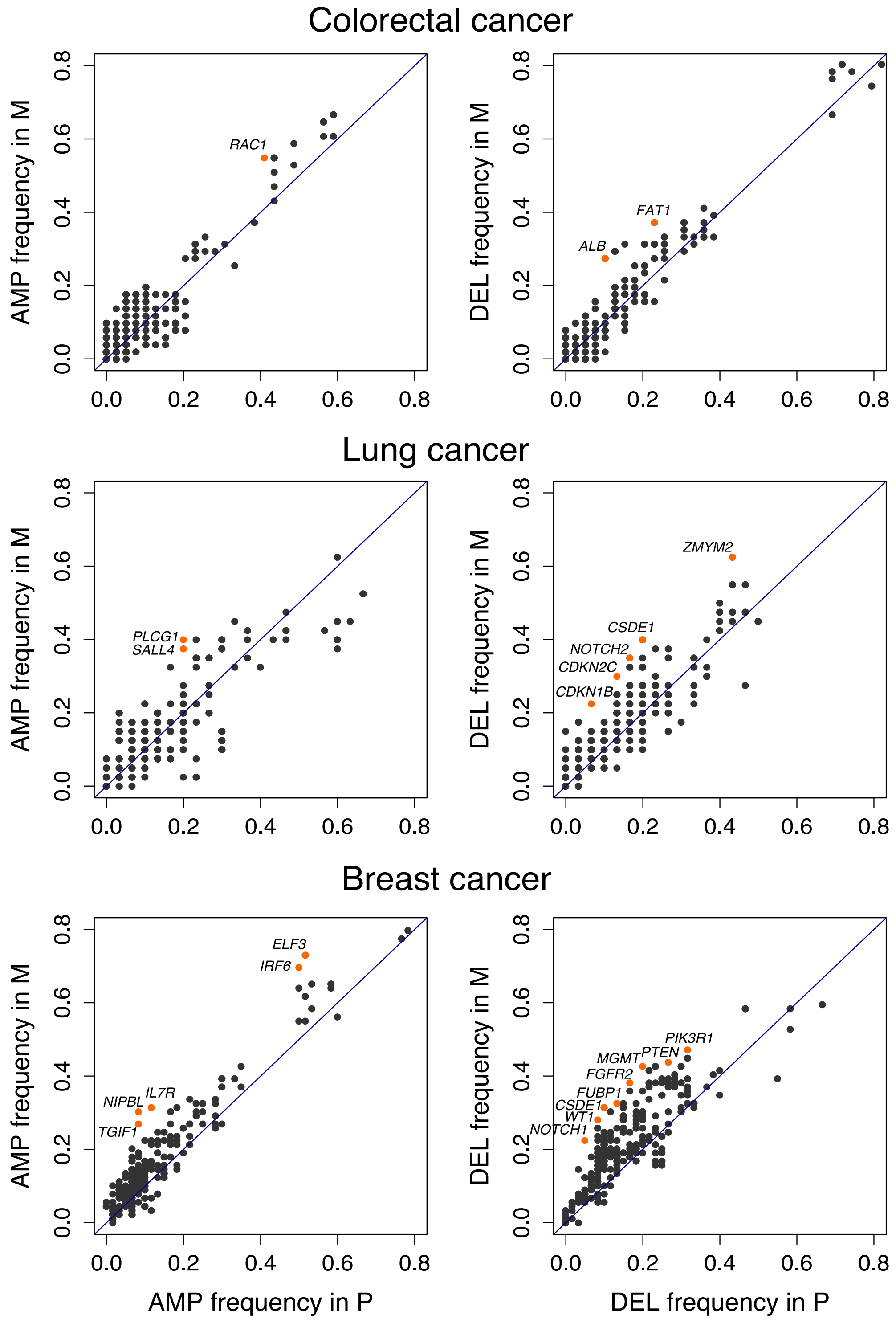

**Supplementary Figure 11. The frequency of SCNAs among putative driver genes in paired primary tumors (P) and metastases (M)**. Left panel, amplifications (AMP) where oncogenes with an increased frequency (≥15%) in the metastasis (M) versus primary (P) are labeled. Right panel, deletions (DEL) where tumor suppressor genes with increased frequency (≥15%) in the metastasis (M) versus primary (P) are labeled.

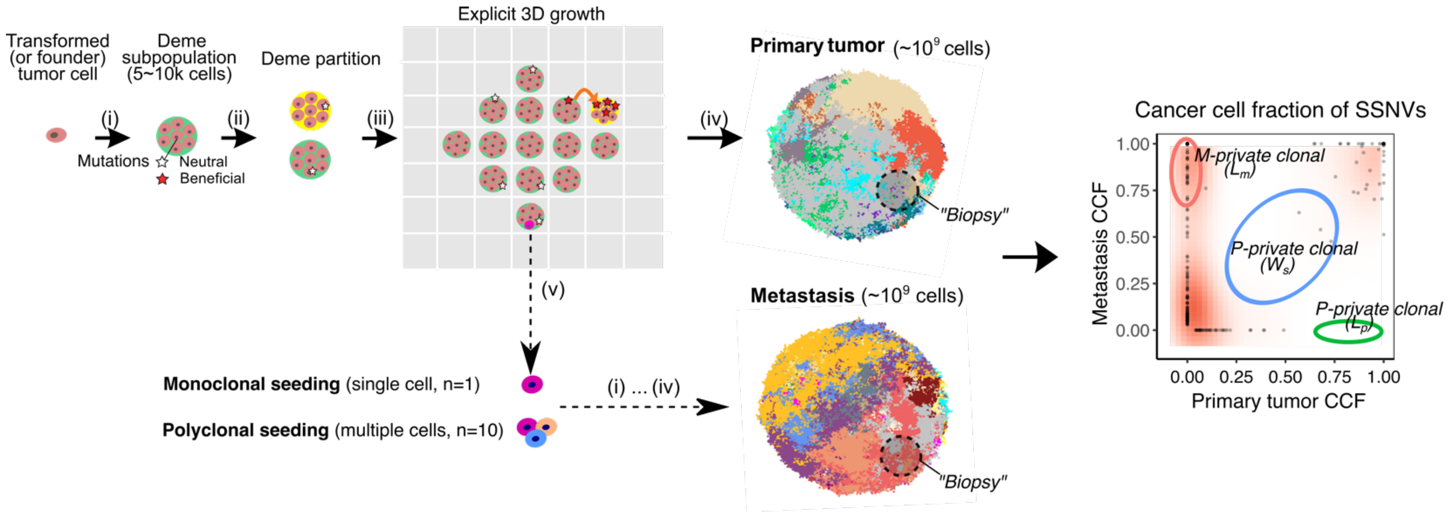

**Supplementary Figure 12. Schematic illustration of a 3-D spatial-agent based model of tumor growth and metastasis.** Tumor growth is simulated via the expansion of deme subpopulations (mimicking the glandular structures often found in epithelial tumors and metastases) within a defined 3-D cubic lattice according to explicit rules dictated by spatial constraints, where cells within each deme are well-mixed and grow via a stochastic branching (birth-death) process, as described in Methods. To model monoclonal seeding, a single cell at the tumor periphery was randomly sampled as the metastasis founder cell. To model polyclonal seeding, a cluster of cells (n=10) were randomly sampled from the whole tumor in order to maximize the clonal diversity within the metastasis founder cells. Metastatic growth follows the same spatial-constraints as the primary and starts from the metastasis founder cell or cell cluster. The final sizes of both the primary tumor and metastasis is ~$10^9$ cells (~$2\times10^5$ demes). Clonal selection is modeled by assuming a constant beneficial mutation rate that alters the cell birth/death probability according to the selection coefficient (denoted *s*). By simulating the acquisition of random mutations (neutral or beneficial), tracing the mutational genealogy of each cell as the tumor expands and subsequently spatially sampling (~$10^6$ cells in each sample) and sequencing the 'final' virtual tumor as is done experimentally after resection or biopsy, we obtain the variant allele frequencies (VAF) and cancer cell fraction (CCF) in both primary tumor and metastasis.

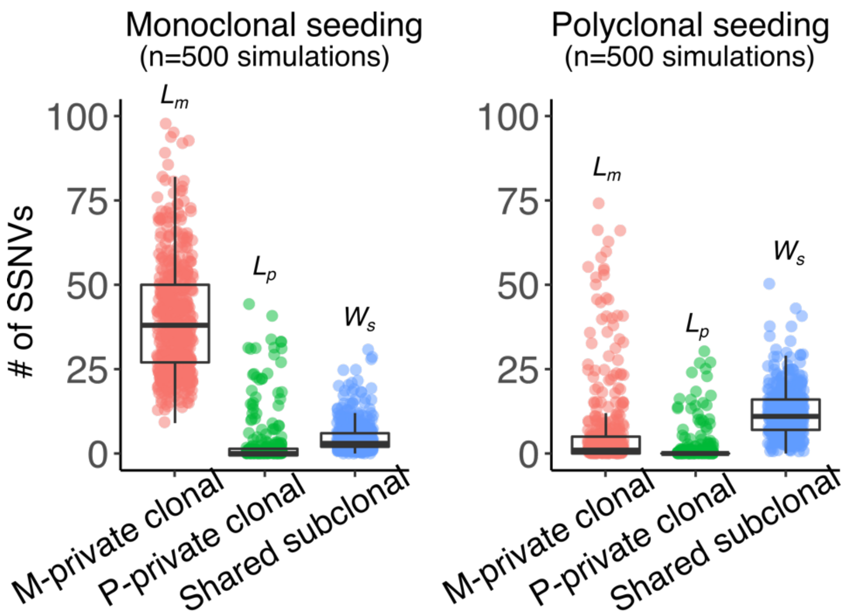

**Supplementary Figure 13. $L_m$, $L_p$ and $W_s$ values in tumors simulated under monoclonal versus polyclonal seeding.** The number of SSNVs in each of the three categories (M-private clonal or $L_m$, P-private clonal or $L_p$, P/M shared subclonal or $W_s$) in the simulated data generated by modeling monoclonal seeding or polyclonal seeding within an agent-based model (Methods) where one sample (~$10^6$ cells) was biopsied from each of primary tumor and metastasis. To model monoclonal seeding, a single cell at the tumor periphery was randomly sampled as the metastasis founder cell. To model polyclonal seeding, a cluster of cells (n=10) were randomly sampled from the whole tumor in order to maximize the clonal diversity within the metastasis founder cells. We employed a mutation rate $u$=0.6 per cell division in the exonic region (corresponding to $10^{-8}$ per site per cell division in the 60Mb diploid coding regions). In order to take account for varying scenarios of tumor growth dynamics, selection and timing of metastatic dissemination, the birth probability $b$ of founding cells, selection coefficient $s$ and primary tumor size at dissemination $N_d$ was randomly sampled from a uniform distribution, $b$~U(0.55, 0.65), log10($s$)~U(-3,-1) and log10($N_d$)~U(4,8), respectively. A total of n=500 virtual P/M pairs were simulated under monoclonal seeding and polyclonal seeding scenarios by randomly sampling these three parameters.

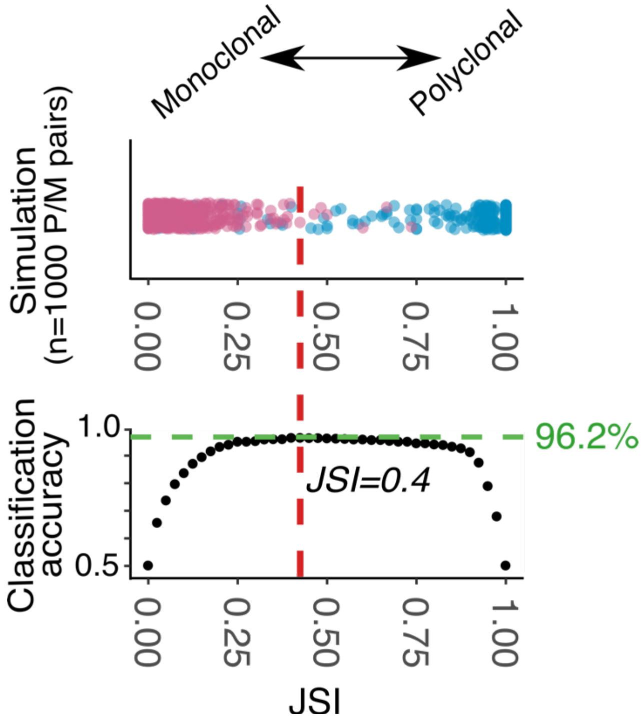

**Supplementary Figure 14. Multi-region sequencing increases the accuracy of classifying monoclonal and polyclonal seeding.** As in Fig. 2c, 1000 virtual P/M tumor pairs were simulated within a spatial tumor growth model with 500 instances of monoclonal seeding (number of metastasis founder cells=1) and 500 instances of polyclonal seeding (number of metastasis founder cells=10). In contrast to Fig. 2c where one "sample" was sequenced, here 4 "samples" (each ~$10^6$ cells) were sequenced from each virtual primary tumor and metastasis pair. The Jaccard Similarity Index (JSI) was computed based on the $L_p$, $L_m$ and $W_s$ from the merged CCFs of four samples in each paired primary tumor and metastases.

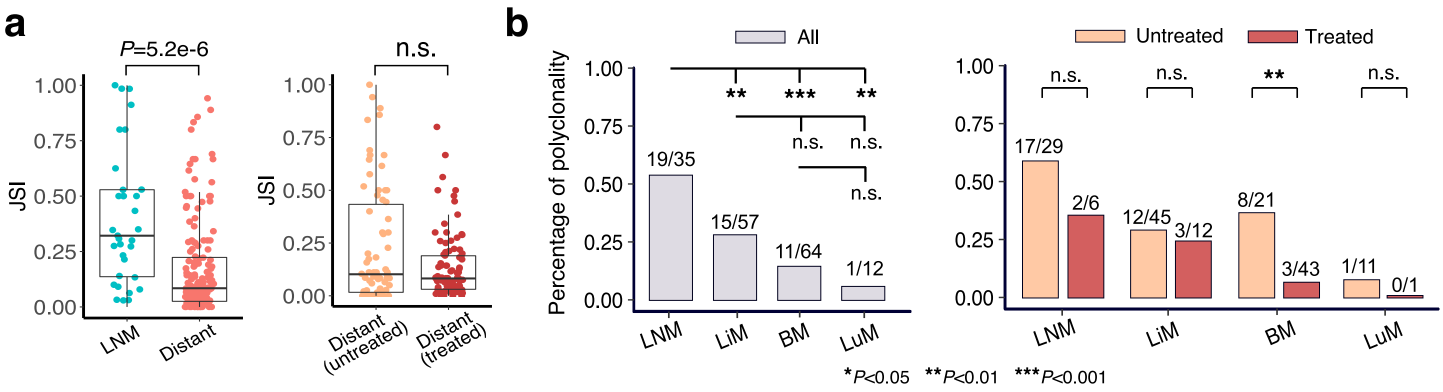

**Supplementary Figure 15. The JSI values in lymph node and distant metastases and the percentage of polyclonal seeding by metastatic sites. (a)** Lymph node metastases (LNM) showed significantly higher JSI than distant metastases. Among distant metastases, untreated metastasis showed higher JSI than treated metastasis although this was not statistically significant. However, using a cutoff of JSI=0.3 to classify polyclonal (JSI≥0.3) versus monoclonal seeding (JSI<0.3), untreated distant metastases showed a significantly higher percentage of polyclonal seeding than treated distant metastases (**Fig. 2e**). *P*-value, Wilcoxon Rank-Sum Test (two-sided). Bar, median; box, 25th to 75th percentile (interquartile range, IQR); vertical line, data within 1.5 times the IQR. (**b**) The percentage of polyclonal seeding among LNM (lymph node metastasis), LiM (liver metastasis), BM (brain metastasis) and LuM (lung metastasis) (left panel) and stratified by treated or untreated (right panel). *P*-value, Fisher's exact test (two sided).

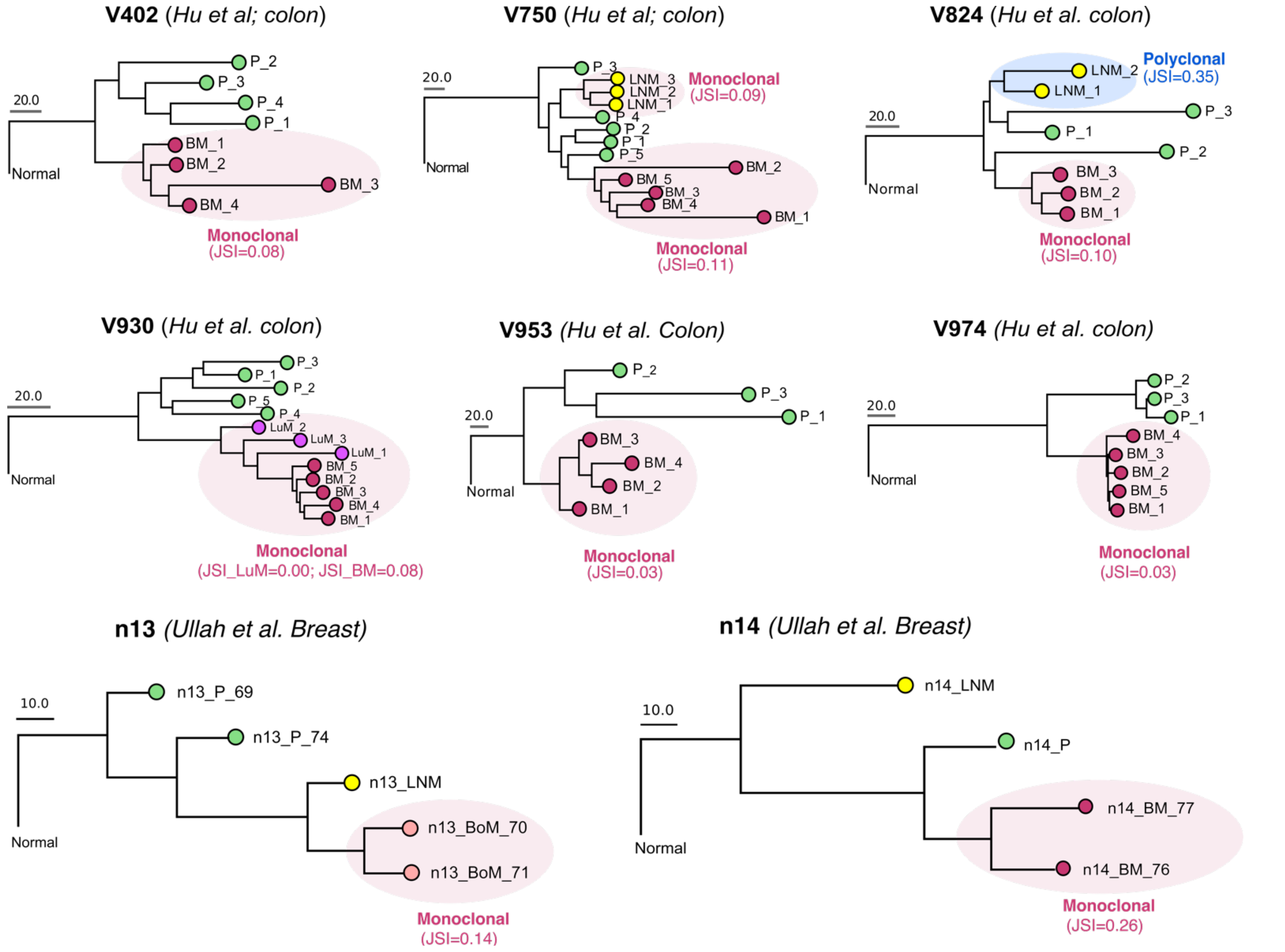

**Supplementary Figure 16. Tumor sample phylogenies based on multi-region sequencing data.** Multi-region sequencing data from six colon cancer patients and two breast cancer patients were used to evaluate tumor phylogenies. P, primary tumor; BM, skin metastasis; LNM, lymph node metastasis; LuM, lung metastasis; BoM, bone metastasis.

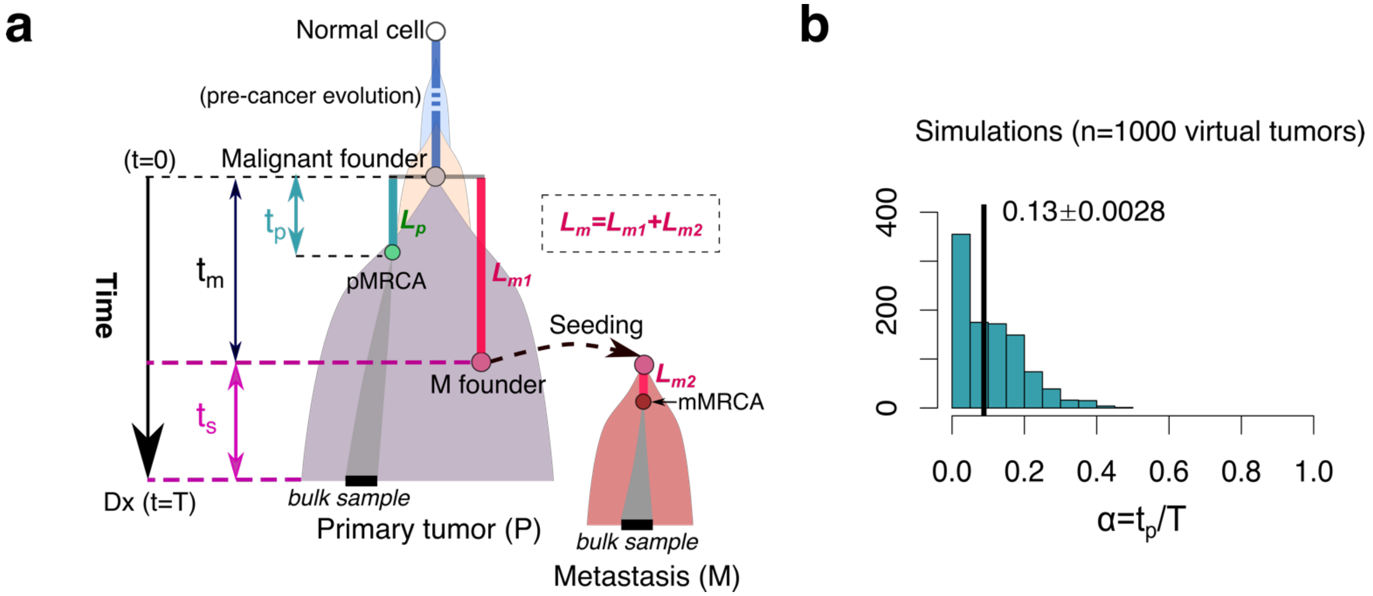

**Supplementary Figure 17. A mathematical method to quantify the chronology of metastatic seeding, $t_s$.** (**a**) Schematic of the parameters used to quantify the metastatic timing $t_s$ in number of years prior to primary tumor diagnosis. We assume metastatic spread occurs at $t_m$ following the emergence of malignant founder lineage for primary carcinoma (a time point denoted as $t$=0). Let $T$ be the time from emergence of malignant founder lineage to diagnosis of the primary tumor (named *primary tumor expansion age*), thus $t_s=T-t_m$. Let $L_p$ and $L_m$ be the number of *private clonal SSNVs in a bulk sample* from paired primary tumor and metastasis, respectively. The clonal mutations defining $L_p$ or $L_m$ are not necessarily clonal in the entire tumor since only a subset of cells are sampled (by biopsy) for sequencing. $L_p$ represents the number of SSNVs that occurred from the emergence of primary tumor founder to *the most recent common ancestor of cell lineages in a bulk sample* (at time denoted pMRCA). This time span is denoted as $t_p$. Similarly, $L_m$ denotes the number of SSNVs occurred from the emergence of primary tumor founder to the most recent common ancestor in a bulk sample from metastasis (at time denoted by mMRCA). $L_m$ includes: (i) the number of M-private clonal mutations that occur within the primary tumor (denoted by $L_{m1}$) and (ii) the number of M-private clonal mutations that occur after cells have disseminated from the primary tumor (denoted by $L_{m2}$), namely $L_m = L_{m1} + L_{m2}$. (**b**) Estimation of $\alpha$ by simulating an agent-based model of spatial tumor evolution (Methods). The mean $\alpha$ and standard deviation from 1000 simulated tumors are shown. Parameter values used in the simulations: neutral mutation rate $u$=0.6 (per cell division in exonic regions); advantageous mutation rate $u_s=1\times10^{-5}$ per cell division and selection coefficient $s$=0.1; birth probability $b$ is sampled from a uniform distribution $b$~U(0.55,0.65) (death probability $d$=1-$b$) to model varying growth rates; final tumor size, $10^9$ cells and biopsy sample size, ~$10^6$ cells; mean sequencing depth, 100X.

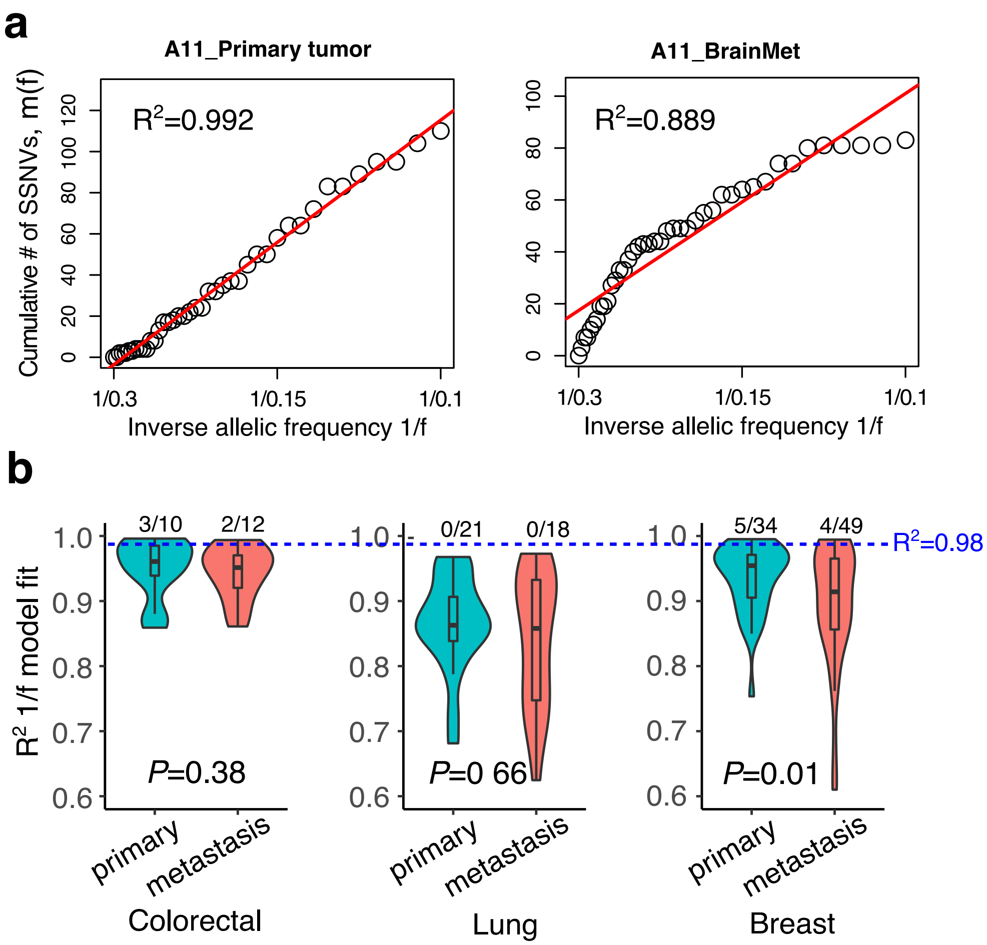

**Supplementary Figure 18**. **Test of neutral tumor evolution by variant allelic frequency (VAF) distribution of subclonal SSNVs.** (a) Representative example, A11 (breast cancer), for the model fitting m(f) ~ f, where m(f) is the number of subclonal SSNVs with VAF larger than f. Both the primary tumor and metastasis are shown. Subclonal SSNVs within the adjusted VAF range 0.1-0.3 were used for this analysis. (b) Violin plot of model fit R2 values in the primary tumor and metastasis, respectively, for the three cancer types. Tumors with R2≥0.98 and R2<0.98 were considered to follow and reject neutral evolution, respectively. The numbers correspond to the number of tumors identified as having patterns consistent with neutral evolution out of the total number of evaluable tumors (those with at least 20 subclonal SSNVs in the range 0.1-0.3 of adjusted VAF). *P*-value, Wilcoxon Rank-Sum Test (two-sided). Bar, median; box, 25th to 75th percentile (interquartile range, IQR); vertical line, data within 1.5 times the IQR.

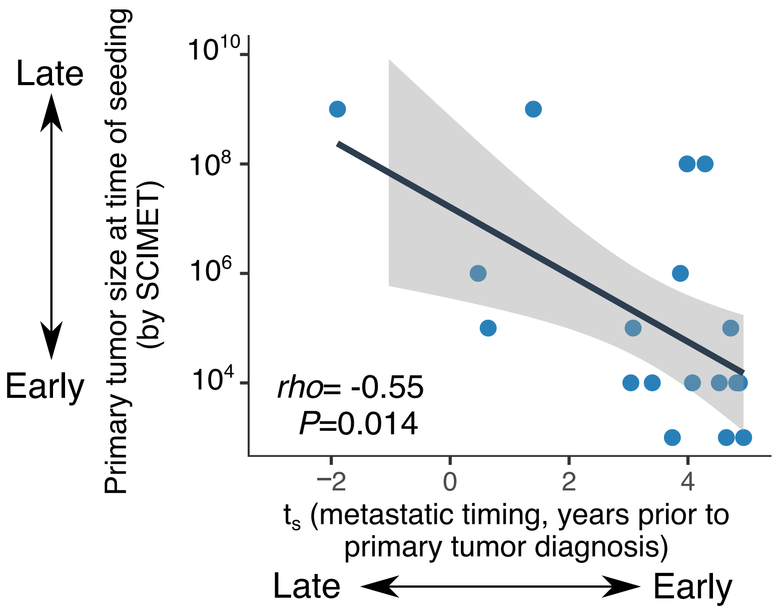

**Supplementary Figure 19**. **Concordance between methods for estimating the timing of metastatic seeding in colorectal cancers**. The x-axis indicates $t_s$ values corresponding to the time of metastatic seeding (years prior to primary tumor diagnosis); the y-axis corresponds to primary tumor size at time of metastatic seeding estimated by SCIMET [16]. SCIMET uses approximate Bayesian computation (ABC) combined with a spatial tumor growth modeling to simulate primary tumor growth and metastatic dissemination at a given primary tumor size (denoted by $N_d$) while $N_d$ values were sampled from a uniform prior (discrete) distribution $\widetilde{N_d}$ ~ {$10^3$, $10^4$, …, $10^9$}. SCIMET yields the posterior probability at each prior $\widetilde{N_d}$ value by fitting a statistical model to the simulated CCF versus the observed CCF values for each P/M pair. $\widetilde{N_d}$ values with the highest posterior probability were reported as the estimated $N_d$. Spearman's correlation (*rho*) and *P*-value are reported. The correlation is negative since here we estimate backward time whereas SCIMET estimates forward time.

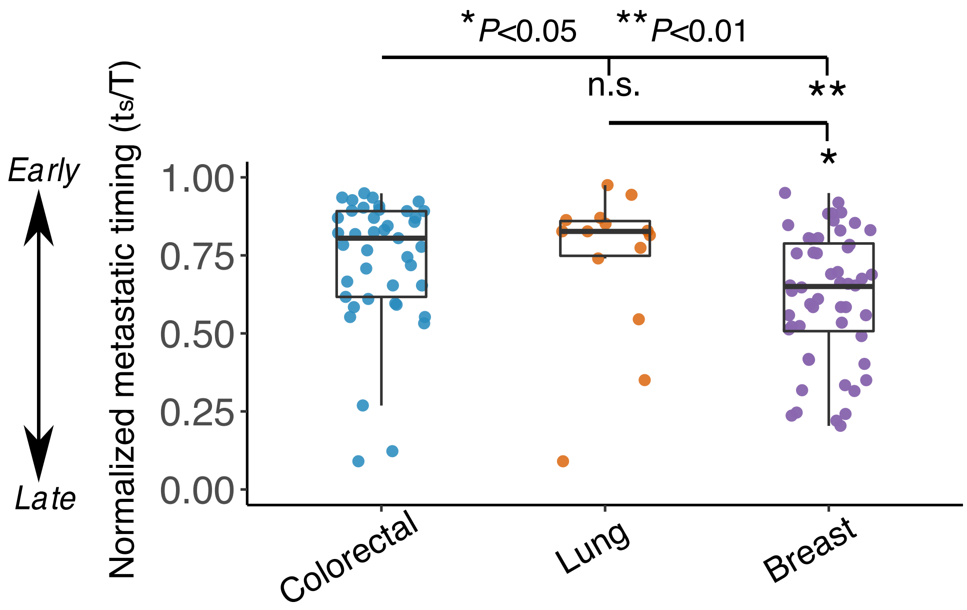

**Supplementary Figure 20. The estimated timing of metastatic seeding normalized to primary tumor age ($t_s/T$).** *P*-value, Wilcoxon Rank-Sum Test (two-sided). Bar, median; box, 25th to 75th percentile (interquartile range, IQR); vertical line, data within 1.5 times the IQR. Only distant monoclonal metastases were considered when estimating the time of metastasis; cases with negative $t_s$ values by Eq(1) were excluded.

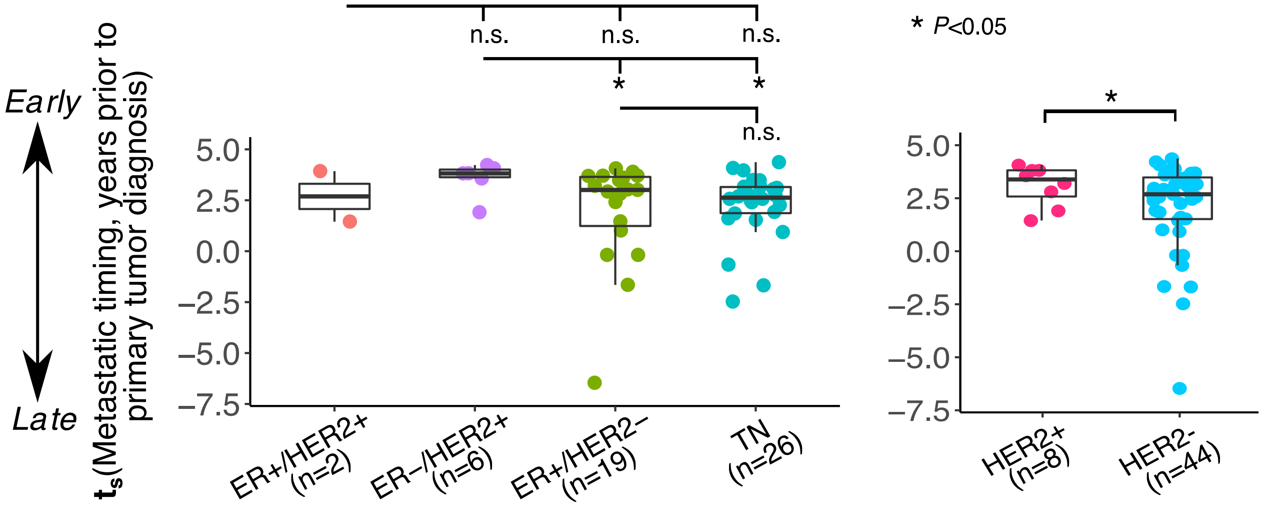

**Supplementary Figure 21. The estimated chronology of metastatic seeding ($t_s$) prior to diagnosis of the primary tumor across the major breast cancer by subtypes.** The four major histopathologic subtypes are considered, namely ER+/HER2+, ER-/HER+, ER2+/HER- and triple negative (TN). In the right panel, HER2+ includes ER+/HER2+ and ER-/HER+ while HER- includes ER2+/HER- and TN. Only distant monoclonal metastases were considered for estimation of the time of dissemination. *P*-value, Wilcoxon Rank-Sum Test (two-sided). Bar, median; box, 25th to 75th percentile (interquartile range, IQR); vertical line, data within 1.5 times the IQR.

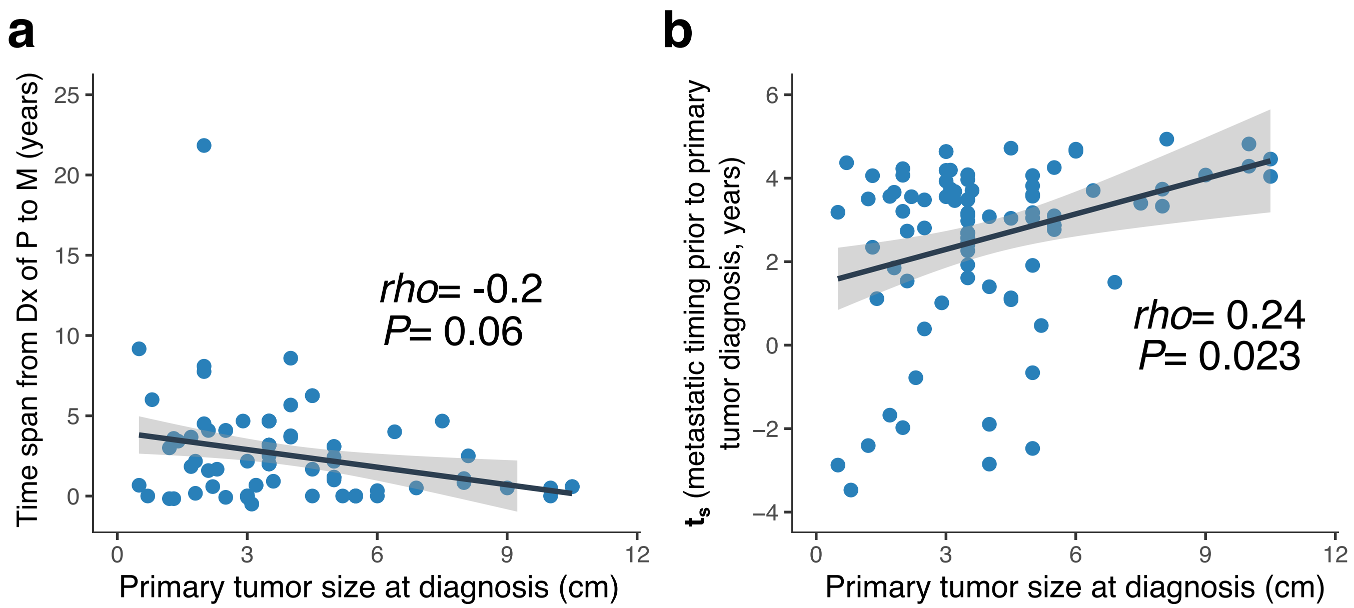

**Supplementary Figure 22. Association of the timing of metastatic seeding with primary tumor size at diagnosis. (a)** Primary tumor size at diagnosis (diameter, cm) is associated with time from diagnosis of the primary tumor to metastasis. (**b**) Primary tumor size at diagnosis (diameter, cm) is associated with the timing of metastatic seeding prior to diagnosis of the primary tumor. A total of n=88 metastases with information on primary tumor size are included. Spearman's correlation (*rho)* and *P*-values are shown.

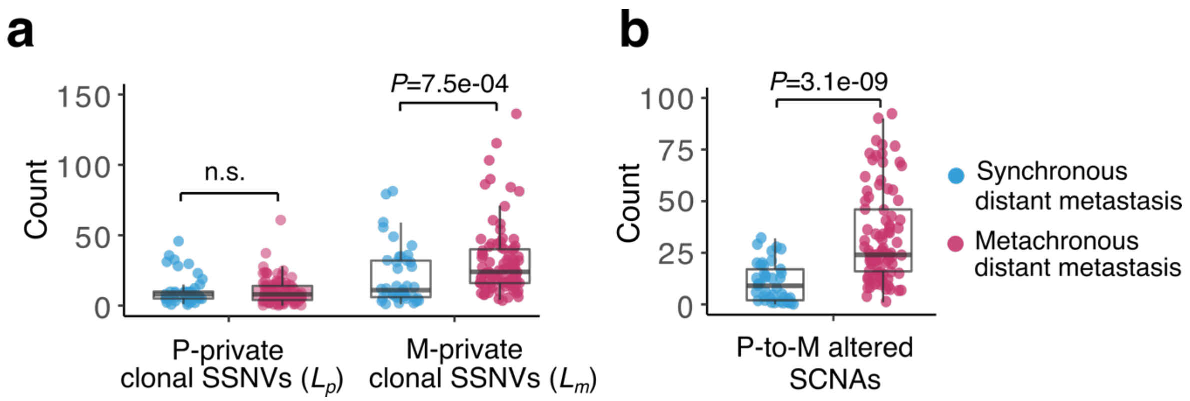

**Supplementary Figure 23. Later metastatic seeding (metachronous metastasis) is associated with higher genomic divergence in the primary tumor.** (**a**) The number of P-private clonal SSNVs and M-private clonal SSNVs in synchronous (distant and monoclonal, n=41) and metachronous (distant and monoclonal, n=80) metastases, respectively. (**b**) The number of P-to-M altered SCNAs in synchronous and metachronous metastases, respectively; *P*-values, two-sided Wilcoxon Rank-Sum Test. Bar, median; box, 25th to 75th percentile (interquartile range, IQR); vertical line, data within 1.5 times the IQR.

### Supplementary Note

#### A mathematical model for estimating the chronology of metastatic seeding

We developed a mathematical method to quantity the chronology of metastatic seeding by leveraging SSNVs as markers of somatic cell divisions. As illustrated in **Fig. S17a**, we assume metastatic spread occurs at $t_m$ following the emergence of malignant founder lineage for primary carcinoma (namely transformation; a time point denoted as $t$=0). Let $L_p$ and $L_m$ be the number of *private clonal SSNVs in a bulk sample* from paired primary tumor and metastasis, respectively. The clonal mutations defining $L_p$ or $L_m$ are not necessarily clonal in the entire tumor since only a subset of cells are sampled (by biopsy) for sequencing. As such, $L_p$ is typically not 0, as can be seen from the patient sequencing data in our cohort (*median of* $L_p$=8, interquartile range or IQR=4−14; **Fig. 23a**). $L_p$ represents the number of SSNVs that occurred from the emergence of primary tumor founder to *the most recent common ancestor of cell lineages in a bulk sample* (at time denoted pMRCA; **Fig. S17a**). This time span is denoted as $t_p$. Similarly, $L_m$ denotes the number of SSNVs occurred from the emergence of primary tumor founder to the most recent common ancestor in a bulk sample from metastasis (a time point denoted by mMRCA; **Fig. S17a**). $L_m$ includes: (i) the number of M-private clonal mutations that occur within the primary tumor (denoted by $L_{m1}$) and (ii) the number of M-private clonal mutations that occur after cells have disseminated from the primary tumor (denoted by $L_{m2}$), namely $L_m = L_{m1} + L_{m2}$. Given the definitions above, we have:

$$\frac{L_m}{L_p} > \frac{L_{m1}}{L_p} = \frac{u_2 t_m}{u_1 t_p} \qquad \textit{(S1)}$$

where $u_1$ and $u_2$ is the mutation rate along the cell lineage in primary tumor bulk sample and metastasis bulk sample, respectively. Given the comparable SSNV burden in primary and metastatic lesions (**Figs. S3-S4**), as also observed recently in a large pan-cancer whole genome sequencing cohort of metastatic tumors [1], we assume a constant mutation rate amongst cell lineages after the emergence of malignant founder cell, such that $u_1 \approx u_2$, as such *Eq.(S1)* reduces to:

$$\frac{L_m}{L_p} > \frac{t_m}{t_p} \qquad \textit{(S2)}$$

Let $T$ be the time from emergence of malignant founder lineage to diagnosis of the primary tumor (named *primary tumor expansion age*) and let $t_s$ be the time of metastatic seeding prior to primary tumor diagnosis, thus:

$$t_s = T - t_m > T - t_p \times \frac{L_m}{L_p} \qquad \textit{(S3)}$$

Denoting $\alpha = t_s/T$, Eq. (S3) can be expressed as:

$$t_s > (1 - \frac{L_m}{L_p}\alpha) \times T \qquad \textit{(S4)}$$

Therefore, $(1-\frac{L_m}{L_p}\alpha)\times T$ is a lower bound of $t_s$ and a conservative estimate of the timing of metastatic seeding. Large $(1-\frac{L_m}{L_p}\alpha)\times T$ indicates early dissemination while small $(1-\frac{L_m}{L_p}\alpha)\times T$ indicates relatively late dissemination. For this reason, $t_s$ can be approximated by:

$$t_s \approx (1-\frac{L_m}{L_p}\alpha)\times T \qquad (S5)$$

Both $\alpha$ and *T* are unknown. $\alpha$ is anticipated to be small since it represents the time during which *private* and *clonal* mutations accrue in the primary tumor sample (**Fig. S17a**), and because bulk sequencing only allows for the detection of high-frequency mutations [2,3]. We applied our established agent-based model of spatial tumor evolution [4,5] to simulate 1000 virtual tumors (each ~$10^9$ cells) with varying growth parameters where the birth probability at each cell generation for the founding cell is *b*~U(0.55, 0.65) (death probability *d*=1-*b*). Here, a bulk sample from a biopsy (~$10^6$ cells, median depth=100X) was sequenced (*in silico*) for each tumor, resulting in $\tilde{\alpha}$=0.13±0.0028 (**Fig. S17b**). Here stringent selection (selection coefficient, s=0.1) was assumed rather than neutral evolution given that for the majority of primary tumors in this cohort (88% or 57/65 evaluable tumors), the allelic frequency distribution was not consistent with neutral evolution (**Fig. S18**).

We use a Gompertzian tumor growth model [6,7] to estimate primary tumor expansion age *T* for the three cancer types in this study. In brief, the number of tumor cells in a Gompertzian growth model can be described as follows:

$$N(t) = N_0 e^{\frac{r}{\beta}(1-e^{-\beta t})} \qquad (S6)$$

where $r$ is the growth rate, $\beta$ is the exponential rate of decrease of the growth rate and $N_0$ the tumor cell number at *t*=0, where $N_0 = 1$. Given the tumor size carrying capacity $K$ (equal to $e^{\frac{r}{\beta}}$), the tumor size doubling time (*DT*) and primary tumor size at diagnosis ($N_p$), $r$ and $\beta$ can be estimated as:

$$\tilde{\beta} = ln\left[\frac{\ln(K) - ln(N_p)}{\ln(K) - ln(2N_p)}\right]/DT$$

$$\tilde{r} = \tilde{\beta} \times ln\,(K) \qquad (S7)$$

The maximal size of human tumors considered here is ~1000 cm(3) [8] which is equivalent to ~$10^{11}$ cells when assuming $10^8$ cells in 1 $cm^3$ tumor tissue [9], thus the carrying capacity is $\tilde{K}$=$10^{11}$. The tumor doubling time *DT* at diagnosis for breast, colon and lung cancers was specified based on a comprehensive literature search for DT in untreated primary tumors (**Table S8**), weighted by the number of patients in each study). The median *DT* (estimate $\widetilde{DT}$) was used in *Eq.(S7).* The average cell number in the primary tumor at diagnosis is $N_p = \frac{4}{3}\pi(D/2)^3 \times 10^8$ where *D* is the median primary tumor size (diameter) at diagnosis and the following values of $\widetilde{D}$ = 4.5 cm, 3.2 cm and 2.0 cm were used for colorectal [10], lung [11] and breast [12] cancers, respectively.

It should be noted that negative values of $t_d$ can occur when the $L_m/L_p$ ratio is large per *Eq.(S5)* under the following scenarios: 1) metastasis was seeded after primary tumor diagnosis and the metastasis seeding cell further accumulated many specific mutations after surgical resection of the primary tumor leading to large $L_m$ values; 2) metastatic seeding before diagnosis of the primary accompanied by a large number of private clonal mutations during metastatic growth (for example due to stringent selection for resistance associated mutations) leading to a large $L_m$. However, bulk sequencing (<100X) has limited power to distinguish whether "M-private" clonal mutations were present but at undetectable frequencies in the primary tumor or whether they occurred after metastatic seeding. Due to this uncertainty, negative estimates of $t_d$ are deemed unreliable and cases where seeding was estimated to occur even after diagnosis of the metastatic relapse were excluded (n=12 for breast, 1 for colorectal and 1 for lung cancer, respectively).

### Supplementary Tables

**Supplementary Table S8.** Reported tumor size doubling time (DT) by cancer type.

| Study | Number of patients | DT (days) |
|---|---|---|
| <u>*Colorectal cancer*</u> | | |
| Bolin, *et al* (1983) [13] | 27 | 195 |
| Choi, *et al* (2012) [14] | 15 | 402 |
| Choi, *et al* (2012) [14] | 2 | 256 |
| Choi, *et al* (2012) [14] | 3 | 219 |
| Choi, *et al* (2012) [14] | 9 | 183 |
| Choi, et al (2012) [14] | 15 | 292 |
| Tata, *et al* (1984) [15] | 11 | 243 |
| Welin, *et al* (1963) [16] | 20 | 620 |
| Umetani, *et al* (2000) [17] | 11 | 204 |
| ***Weighted median*** | | **233 (IQR, 191–341)** |
| <u>*Lung cancer*</u> | | |
| Arai, *et al* (1994) [18] | 237 | 166 |
| Garland, *et al* (1963) [19] | 41 | 162 |
| Geddes (1979) [20] | 228 | 102 |
| Jennings, *et al* (2006) [21] | 149 | 161 |
| Mizuno, *et al* (1984) [22] | 50 | 136 |
| Schwartz (1961) [23] | 13 | 78 |
| Spratt, *et al* (1963) [24] | 22 | 112 |
| Spratt and Spratt (1964) [25] | 34 | 88 |
| Usuda, *et a*l (1994) [26] | 165 | 164 |
| Weiss (1974) [27] | 28 | 183 |
| ***Weighted median*** | | **159 (IQR, 104–212)** |
| <u>*Breast cancer*</u> | | |
| Fornvik, *et al* (2015) [28] | 31 | 282 |
| Galante, *et al* (1981) [29] | 196 | 60 |
| Heuser, *et al* (1979) [30] | 32 | 268 |
| Kuroishi, *et al* (1990) [31] | 118 | 174 |
| Kusama, *et al* (1972) [32] | 199 | 105 |
| Lundgren (1977) [33] | 15 | 211 |
| Peer, *et al* (1993) [34] | 46 | 80 |
| Peer, *et al* (1993) [34] | 188 | 157 |
| Peer, *et al* (1993) [34] | 55 | 188 |
| Ryu, *et al* (2014) [35] | 37 | 241 |
| Ryu, *et al* (2014) [35] | 12 | 162 |
| Ryu, *et al* (2014) [35] | 17 | 103 |
| Tabbane, *et al* (1989) [36] | 75 | 115 |
| von Fournier, *et al* (1980) [37] | 147 | 212 |
| von Fournier, *et al* (1985) [38] | 200 | 220 |
| ***Weighted median*** | | **149 (IQR, 100–164)** |

**Supplementary Table S9**. Parameters associated with metastatic seeding times.

| Cancer type | $\widetilde{D}$ (cm) | $\widetilde{N_p}$ (cells) | $\widetilde{K}$ (cells) | $\widetilde{DT}$ (days) | $\tilde{r}$ | $\tilde{\beta}$ | $\tilde{T}$ *(years)* | $\tilde{\alpha}$ |
|---|---|---|---|---|---|---|---|---|
| Colorectal | 4.5 | $4.77 \times 10^9$ | $1 \times 10^{11}$ | 233 (191–341) | 0.028 (0.019–0.034) | 0.0011 (0.0008–0.0014) | 5.2 (4.3–7.7) | 0.13 |
| Lung | 3.2 | $1.72 \times 10^9$ | $1 \times 10^{11}$ | 159 (104–212) | 0.030 (0.029–0.047) | 0.0012 (0.0011–0.0019) | 4.3 (2.7–4.4) | 0.13 |
| Breast | 2.0 | $4.18 \times 10^8$ | $1 \times 10^{11}$ | 149 (100–164) | 0.023 (0.016–0.033) | 0.0009 (0.0006–0.0013) | 4.6 (3.2–6.6) | 0.13 |
